# Principles of protein abundance regulation across single cells in a mammalian tissue

**DOI:** 10.1101/2025.09.17.676955

**Authors:** Andrew Leduc, Gergana Shipkovenska, Yanxin Xu, Christina Jiang, Alexander Franks, Nikolai Slavov

## Abstract

Protein synthesis and clearance are major regulatory steps of gene expression, but their in vivo regulatory roles across the cells comprising complex tissues remains unexplored. Here, we systematically quantify protein synthesis and clearance across over 4,200 cells from a primary tissue. Through integration with single-cell transcriptomics, we report the first quantitative analysis of how individual cell types regulate their proteomes across the continuum of gene expression. Our analysis quantifies the relative contributions of RNA abundance, translation, and protein clearance to the abundance variation of thousands of proteins. These results reveal a putative organizing principle: The contributions of both translation and protein clearance are linearly dependent on the cell growth rate. Further, we find that some proteins are primarily regulated by one mechanism (RNA abundance, translation, or clearance) across all cell types while the dominant regulation of other proteins is cell-type specific. Age related changes in protein abundance are cell-type specific and correlated to changes in protein clearance. Our reliable multimodal measurements enabled quantifying and functionally interpreting molecular variation across single cells from the same cell type. The protein-protein correlations are substantially stronger than the mRNA-mRNA ones both for directly interacting proteins and for functional protein sets. This difference is mediated by protein clearance regulation. Further, the protein correlations allow identifying cell-type specific functional clusters. These clusters vary across cell types, revealing differences in metabolic processes coordination, partially regulated by protein degradation. Our approach reveals organizing principles determining the relative contributions of translation and protein clearance and provides a scalable framework for investigating protein regulation in mammalian tissues.

## Main

For decades, protein abundance has been approximated by mRNA abundance because direct protein measurements were technically difficult^1,2^. Now, technological progress^3–6^ has enabled numerous proteomic studies across diseases and aging^7^, establishing many instances where mRNA levels are poorly predictive of protein levels^1,2,8^. Despite this progress, the mechanisms regulating protein abundance that contribute to the observed discordance with mRNA remain understudied.

Proteostasis has been studied extensively using cell lines in bulk tissues, and these studies helped establish the core kinetic and computational frameworks for turnover analysis^9–13^. Further, these studies have revealed global trends in protein degradation^14^ and regulatory mechanisms of protein degradation, including ubiquitin ligase substrates^15,16^. However, cultured cell lines lack many of the physiological inputs present in an intact tissue, such as cell-cell signaling, extracellular matrix, immune surveillance, and organismal-level processes, such as autophagy induction and tissue turnover and regeneration. Furthermore, prior work has shown that the range and distribution of protein half-lives measured in cell lines can differ substantially from those measured in vivo^12,17^. However, the proteostasis studies of primary tissues so far have been limited to bulk samples reporting population averages of highly specialized cells. The single-cell studies of gene expression regulation within primary tissues have been confined to the regulation of RNA abundance as it can be measured by scalable single-cell RNA sequencing technologies. Here, we extend it to the full path of gene expression, focusing for the first time on the regulation of protein synthesis and clearance in single cells from a complex mouse tissue.

Regulation of protein synthesis and clearance may also contribute to functional differences across individual single cells within a cell type. For example, a subset of basal stem cells within the tracheal epithelium have been identified as injury-resistant reservoirs with heightened proliferative capacity^18^. Since synthesis and clearance scale with growth rate^12,19,20^, variation in proliferative state may shape protein abundances across basal cells not reflected at the level of mRNA abundance. Comparable functional heterogeneity has been observed even in the absence of clearly defined subpopulations, such as the variable differentiation of isogenic iPSCs exposed to the same stimulus^21^ or the ability of a subset of cancer cells to survive drug treatment^22,23^. The extent to which such functional heterogeneity is a general feature of cells within a defined type, however, remains unclear.

A central obstacle to addressing this question with single-cell mRNA sequencing is that the sparse and stochastic nature of these measurements typically restricts analysis to comparisons between clusters of cells rather than continuous variation within a cluster^21,24,25^. Thus, and mRNA differences within a cell type are often difficult to interpret functionally^26^. Proteins offer a more direct readout of functional state as they are the effectors of most cellular processes. Further, their higher copy number and longer half-lives make measurements substantially less stochastic than mRNA^9,27^. Covariation across multiple proteins may then provide even stronger evidence of shared functional state, and may be more readily detected at the protein than at the mRNA level. Jointly analyzing protein abundance and clearance covariation across single cells within a cell type may both identify cell-type-specific functional variation^28^ and reveal how the protein expression underlying it is regulated.

Using scalable single-cell mass spectrometry (MS), we sought to elucidate principles of protein abundance regulation across the full continuum of gene expression. To quantify the contributions of RNA abundance, translation and protein clearance across diverse cell types, we chose murine trachea as a model, Fig. 1a. We found the tracheal epithelium to be a good model system for our analysis as it contained both a variety of cell types for contrasting cell-type specific protein regulation as well as heterogeneous stem epithelial cells ideal for within cell type analysis^18,29^.

Our analysis uncovers the ways in which the primary mode of gene regulation differs across individual proteins and functional groups of proteins. Further, the protein abundance regulation varies across cell types, even for the same proteins. Single-cell level analysis reveals substantial protein covariation across single cells of the same type, reflecting cell-type specific functional protein coordination. Dissecting this regulation highlights the role of protein clearance; it shapes both protein abundance and protein-protein covariation in individual single cells, substantially contributing to discordance between mRNA and protein covariation patterns.

## Results

### In vivo metabolic labeling enables quantification of regulation in single cells

To enable this analysis, we introduced combined multiplexing of isotopic peaks from metabolic pulse labeling with isotopic chemical barcoding for sample multiplexing via the plexDIA framework^30^, Fig. 1b. Previously, plexDIA has been used for sample multiplexing (enabling measuring thousands of cells from a human tissue^31^) and for metabolic pulse labeling for protein turnover measurements^32,33^, but not for both simultaneously. We reasoned that combining these different approaches could enable quantifying protein synthesis and clearance rates while increasing the throughput of single cell analysis, Fig. 1b. We achieved this by a metabolic pulse with +6 Da heavy lysine combined by multiplexing single cells with mTRAQ Δ0 and Δ8 tags creating a 16 Da offset between lysine containing peptides from different single cells. To prepare single cell samples, we leveraged the nPOP glass-slide sample preparation^34,35^. This approach facilitated preparing 1,300 single cells per batch with multiplexing two cells per set.

This design empowered the analysis of over 4,200 single cells from 6 mice fed with heavy lysine food for either 5 or 10 days, Fig. 1a,c. We aligned this data set with over 36,000 single cells from mice in the space of 2,201 shared gene products quantified at the mRNA and protein level^18^. Mice labeled for 5 days were either 4 months or 24 months old. Each mouse labeled for 10 days was 4 months old as the two 24 month old mice perished on the 9th day of labeling and were excluded from the study. The mRNA sequencing data set consisted of 4 mice, two 4 month and 24 months of age.

We started our analysis by first clustering and annotating each dataset separately based on abundance of common marker proteins and then aligned the cell types, Extended Data Fig. 1a. To validate cell type assignment across data sets, we correlated relative levels of protein and mRNA abundance across cell types. We observed remarkably strong agreement for matched cell types. The strong correlations among independently annotated cell type clusters indicate that our data support robust cell type assignment without the need for more complex integration approaches, Fig. 1d and Extended Data Fig. 1b.

To improve the quantification, we devised strategies to account for artifacts such as variability in liquid chromatography (LC) and missing values. These approaches were added as functionality to our QuantQC package used for downstream processing of searched single cell data^35^. First, due to the large number of LC-MS/MS runs, we noticed the intensities of certain peptides drifting as a function of the run number, Fig. 2a. Similar to previous approaches applied in metabolics^36^, we regressed out unwanted variation for each peptide using a spline fit to the LC run order, see methods. To account for the influence of missing data, we devised a missingness-aware approach to improve fold change estimates across cell types. For each protein, we identified the percentage of cells with missing values caused by either biological or technical factors^37^, allowing us to scale fold changes proportionally to biological missingness, see methods. Technical factors included cell size, sample preparation batch, multiplexing label, and order in which the samples were run on the MS. We identified cell size as a leading factor for technical missingness with LC batch and label explaining missingness to a lesser extent Extended Data Fig. 2b-e. The approach improved agreement between relative protein and RNA abundance estimates (Extended Data Fig. 2f-h), suggesting improved quantification. It should be noted that this correction was only applied for analysis of relative protein levels across cell types and not for regulatory analysis that involves within cell type analysis or at the single cell level.

**Table 1.**
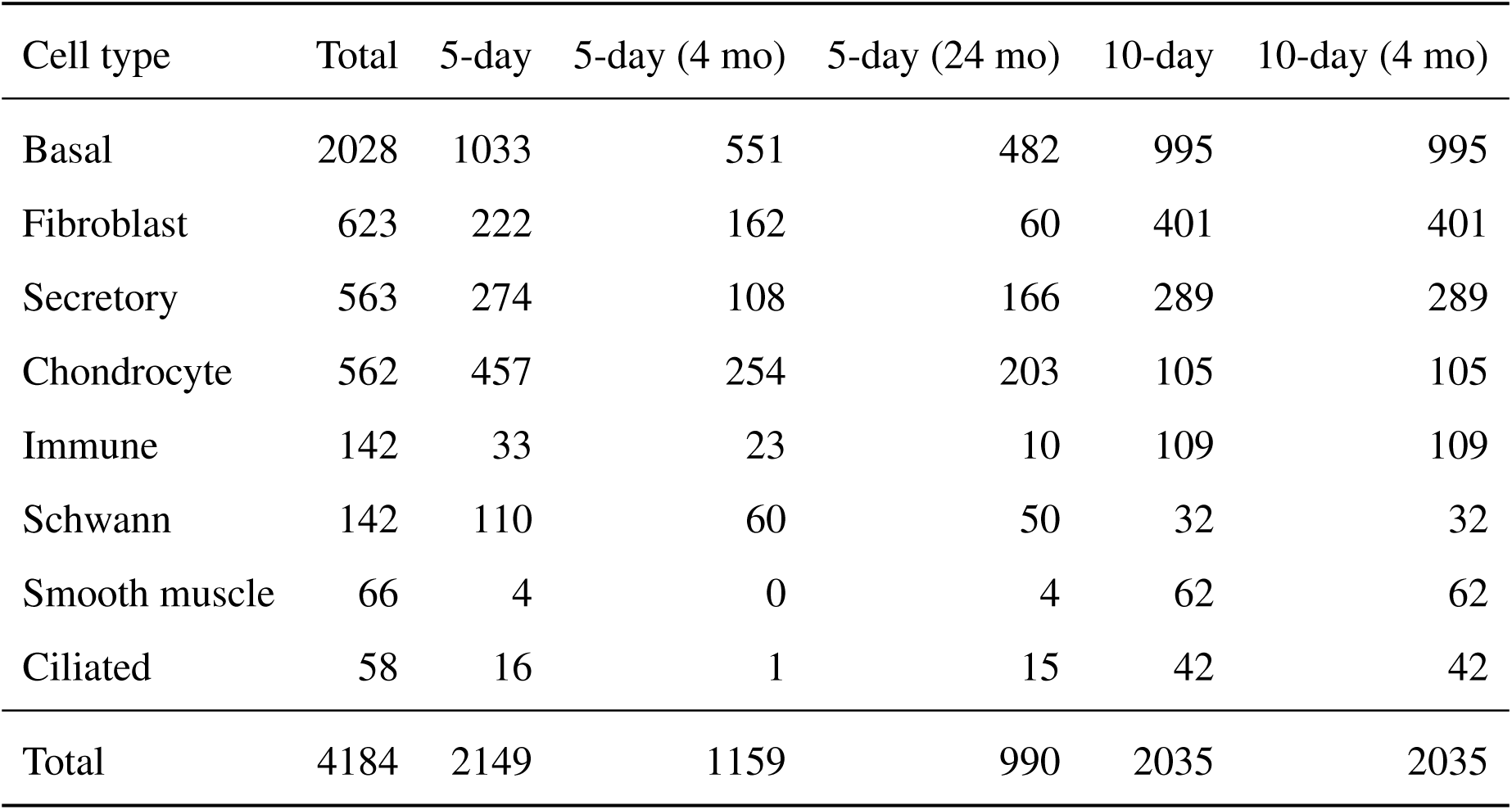
Single-cell counts per cell type, broken out by pulse duration and age. Totals reflect all quantified single cells passing quality control. The 10-day cohort contains only young (4-month) mice, as the two 24-month old mice perished before tissue collection on day 9 of the labeling period.

After controlling for technical artifacts, we then set out to estimate protein clearance rates at the level of cell types and single cells. Accurate estimation of these rates from *in vivo* metabolic labeling depends on both accurate modeling of amino acid recycling^38^, and validating our modeling choices, namely that single cells are approximately at steady state, Extended Data Fig. 3a. First, we computed the exchange of free lysine amino acids in each cell type utilizing miss-cleaved peptides^10^, see methods. The rate of exchange differed depending on cell type, generally higher for epithelial and immune cell types compared to mesenchymal and smooth muscle, Extended Data Fig. 3b. Cell-type specific lysine exchange rates were then used to correct half lives for all single cells across each respective cell type. Correcting for recycling reduced half-life of protein clearance from roughly 7 days on average to 3 days, Extended Data Fig. 3c. Further, applying this correction harmonized clearance rates computed from 5 and 10 day cells, Extended Data Fig. 3d. Because the 10 day old mice cohort perished before tissue collection, we assumed the free heavy amino acid pool percentage for the 10 day mice increased by the same fraction as the light, Extended Data Fig. 3b. Though we explicitly flag this as a limitation, we observed good alignment between the slope of clearance rates from old and young mice, Extended Data Fig. 3e. The small offset may suggest a reduction in overall clearance rates with age^39^.

**Figure 1.**
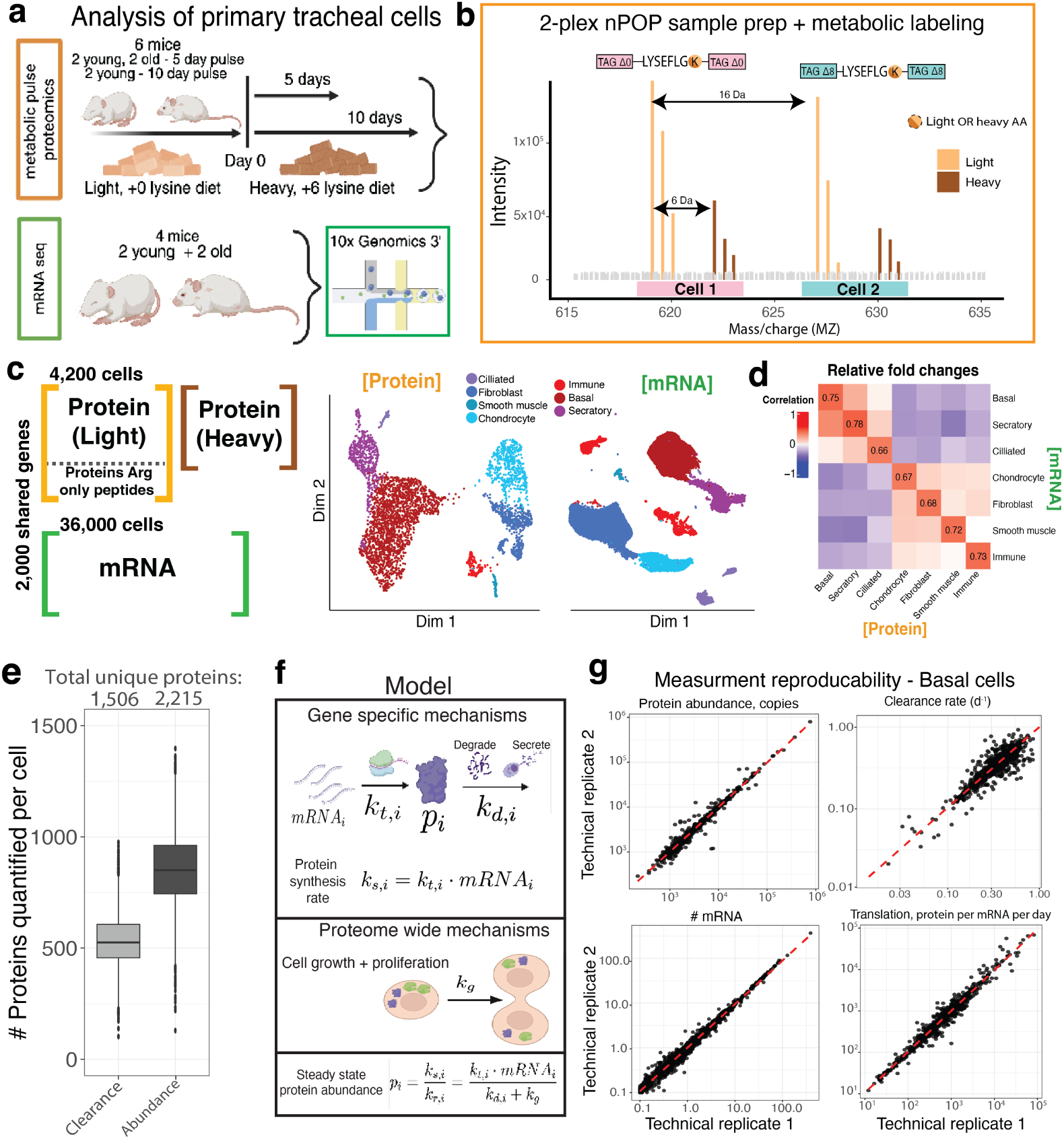
Single-cell metabolic pulse proteomics and transcriptomics in murine trachea. **a**, Mice for scproteomics were fed a diet of +6 Da heavy lysine for 5 days (*n* = 4) or 10 days (*n* = 2). Single cells prepared via nPOP multiplexed sample preparation were labeled with either mTRAQ Δ0 or Δ8. **b**, Mass spectra from precursor ions for peptide LYSEFLGK showing the 4 co-eluting isotopic states. Each of the two multiplexed single cells contributes two peaks: one from unlabeled (+0 Da) lysine and one from metabolically incorporated heavy (+6 Da) lysine. The mTRAQ Δ0 and Δ8 chemical labels impose a 16 Da offset between the two cells, yielding four resolvable precursor masses per peptide. Single cells from the trachea of 4 different mice were processed for sc-mRNA seq by 10x Genomics. **c**, The resulting dataset contains 4,200 single cells with proteomic measurements of peptides containing light and heavy lys and 36,000 cells with RNA data. UMAP visualization shows the corresponding cell types clustered in the space of either protein or RNA abundance. **d**, The relative abundance of protein and mRNA levels in corresponding cell types are strongly correlated (diagonal). **e**, Number of proteins quantified per cell. Across all cells, protein abundances were quantified for 2,215 unique proteins and synthesis and degradation rates for 1,506 unique proteins. Details shown in Extended Data Fig. 6 **f**, Protein abundance at steady-state for a given protein can be modeled as the rate of protein synthesis (which equals mRNA concentration multiplied by translation efficiency, the number of proteins transcribed per mRNA),and the rate of proteins cleared via degradation, secretion, or dilution due to cell growth. **g**, The reproducibility of estimates for protein and mRNA abundance, protein clearance, and translation per mRNA is visualized in basal cells by comparing technical replicates of averaged non-overlapping subsets of single cell.

Since differences in amino acid recycling were identified across ages and cell types, we sought to determine whether significant recycling variation exists across single cells within a cell type. One possible source of such variation might be that more metabolically active cells recycle amino acids faster. We tested this by correlating the per-cell recycling estimate with the average clearance rate across all proteins within basal cells, Extended Data Fig. 3f. We observed no relationship, indicating that recycling estimates were not systematically higher for cells with faster turnover.

To test more broadly for recycling variation caused by other factors, we exploited the fact that each mis-cleaved peptide provides an independent recycling estimate. If cells vary in a coordinated way in their recycling, estimates from different peptides should correlate positively across cells. When we correlated recycling estimates from different peptides across single cells, we observed no correlation on average, Extended Data Fig. 3f, arguing against coordinated differential recycling across single cells. One caveat to this reasoning is that individual proteins might exhibit variable recycling dynamics that would not produce a coordinated across-peptide signal. To evaluate this possibility, we compared the similarity of recycling estimates between peptides mapping to the same protein and peptides mapping to different proteins within a cell, Extended Data Fig. 3g. We observed no statistically significant difference between these two groups, arguing against protein-specific recycling variation as a hidden confound.

While we did not detect evidence for significant recycling variation across single cells, we acknowledge that recycling estimates derived from a limited number of mis-cleaved peptides per cell may be suboptimal for detecting subtle differences. Future work using proteases that generate more peptides with multiple lysines, such as Arg-C, could enable a deeper exploration of single-cell recycling heterogeneity.

Overall, we quantified clearance rates for 1,506 proteins with an average of 514 proteins per cell, Fig. 1e. As clearance rates have previously been shown to vary across cell types cultured in vitro^40^, here we focused on quantifying this variation across the cell types of a tissue. To examine this, we first compare the distribution of clearance rates for each cell type. On average, clearance rates were lowest for chondrocytes and smooth muscle cells while being highest for basal cells, Extended Data Fig. 3h. We then correlated clearance rates across cell types. The average similarity across cell types is *ρ* = 0.50 Extended Data Fig. 3i, much lower than the reliability estimated from independent subsets of cells of the same type, as shown on the diagonal. For example, a correlation of *ρ* = 0.51 between basal cells and chondrocytes while internal estimates for each cell type are above *ρ* = 0.75. This difference strongly suggests cell-type specific rates of protein clearance.

However, it is possible for these measurements to be internally consistent but inaccurate if our steady state assumption is not well justified. To further test the steady-state assumption, we asked whether cells actively transitioning between states. Such transition would violate the steady-state assumption and can be detected in the data. Basal-to-secretory differentiation is a well-established transition in the tracheal epithelium and thus provides a natural test case^29^. We projected basal and secretory cells into a low-dimensional space via principal component analysis and fit a principal curve with 3 degrees of freedom to define a putative differentiation trajectory, Extended Data Fig. 4a. Along this trajectory, the secretory marker Arg2 gradually increased and the basal marker Krt5 gradually decreased, as expected, Extended Data Fig. 4b.

If our system captured the dynamic transition, we would expect to observe an increase in newly synthesized heavy protein lead the light Arg2. This would reflect cells that occupied a basal state with low Arg2 before the pulse and began to synthesize new heavy Arg2 as they started their transition after the pulse, Extended Data Fig. 4c, as previously demonstrated in single-cell metabolic labeling of transcripts^41^. Instead, heavy and light abundance shifted at the same rate for both marker proteins, and the same pattern held for the top 20 varying proteins along the trajectory, Extended Data Fig. 4d.

To test for localized transitions that a whole-trajectory view might miss, we performed a sliding-window analysis analogous to trajectory-lag comparisons used to infer directionality in single-cell RNA velocity^42^. For each position along the trajectory, we defined a reference window of heavy protein abundances at that position and correlated it against light protein profiles at each preceding and following positions, Extended Data Fig. 4d,e. A dynamic transition would produce higher correlations with future light profiles (cells’ current newly synthesized proteome predicts future mature proteome) than with past heavy profiles. We quantified this asymmetry as *θ* = CF−CP, the difference between the average correlation to the three following heavy windows (CF) and the three preceding ones (CP). Across the trajectory, *θ* averaged 0.01 with no sustained positive regions, Extended Data Fig. 4f, indicating that our pulse did not capture a dynamic basal-to-secretory transition. This suggests that the time scales of protein synthesis and degradation are sufficient different from the time scale of basal-to-secretory cell differentiation; this difference supports time scale separation consistent with quasi steady-state modeling and analyses.

To further quantify the reliability of our clearance estimates, we cross-referenced them against two orthogonal comparisons. First, in addition to computing measurement reliability across proteins by comparing estimates from independent subsets of cells within a cell type, we compared estimates for each protein derived from different peptides mapping to it, Extended Data Fig. 5a-c. These peptides provide clearance estimates that are independent of many MS biases such as interferences and ionization^43^. We first examined the raw data by selecting a representative set of 10 proteins spanning the distribution of clearance rates in basal cells, each with at least 4 mapped peptides. Protein identity explained the majority of clearance variation across peptides, Pearson *r* = 0.95, indicating the ability to distinguish protein-specific clearance rates, Extended Data Fig. 5a. To broaden the comparison to all 863 proteins with multiple peptide measurements, we randomly partitioned peptides from each protein into two sets and computed clearance rates from each set independently, averaged across basal cells, Extended Data Fig. 5b. We again observed strong agreement between the two partitions, Pearson *r* = 0.78. This approach likely underestimates the true reliability, since halving the number of peptides per protein increases the noise in each partition’s estimate. Similar agreement was observed within each cell type, Extended Data Fig. 5c.

Second, we compared our estimates to prior work in a similar cell type from Schwanhausser et al.^9^, Extended Data Fig. 5d. That study estimated protein half-lives in cultured NIH3T3 fibroblasts using 10 metabolic pulse timepoints, compared to the two used here. Our tracheal fibroblasts and their NIH3T3 fibroblasts showed a Spearman correlation of *r* = 0.50. Despite the biological differences between fibroblasts cultured in vitro and our in vivo tracheal fibroblasts, this level of agreement falls within the range of cross-study comparisons reported in recent surveys of protein turnover measurements^44^.

Having validated the accuracy of our clearance estimates, we sought to quantify and understand how such regulated protein clearance plays into the larger picture of gene expression. To this end, we inferred a range of parameters spanning the gene expression process. Under steady state, protein synthesis equals the product of protein clearance rate and abundance^12,19^, Fig. 1f. Protein synthesis itself is the joint contribution of the amount of mRNA molecules and the rate at which each molecule is translated into proteins. Our data enabled us to estimate each of these quantities and their influence at the cell type level.

We next sought to examine the reliability of the inferred quantities. For the scRNA-seq, we achieved depth of at least 3,000 unique transcripts per with at least 10k unique molecular identifiers (UMIs) per single cell, comparable to the best published 10x genomics data sets, Extended Data Fig. 6a. For the single-cell proteomics data, we quantified abundances for 2,205 proteins with an average of 805 per single cell, Fig. 1e. Variation in the number of proteins quantified can be attributed to cell size variation, Extended Data Fig. 6b. Further, the strong correlation between cell size and total protein highlights the consistency and precision of the sample preparation. To evaluate the accuracy of protein quantification, we examined the consistency of relative abundances for peptides mapping to the same protein^43^, Extended Data Fig. 6c. For example, the 5 most abundant peptides from the protein Tgm2 show high correlation across single cells, *ρ* = 0.88. On average, peptides originating from a protein correlated with *ρ* = 0.50. As expected, proteins that varied more significantly from cell to cell displayed higher correlations, Extended Data Fig. 6d.

To evaluate the consistency of the estimates within cell types, we partitioned cells into two populations and computed the average mRNA and protein abundance, protein clearance rate and translation rate, Fig. 1g and Extended Data Fig. 6e. The reliability of agreement between the estimates was above *ρ* = 0.85 for protein abundance, mRNA abundance, and translation. Lower correlation, on average *ρ* = 0.69, for clearance rates might reflect the smaller dynamic range of the clearance rates. These estimates allowed us to account for the influence of measurement noise when determining how different regulatory modes impact protein abundance across cell types.

### Growth rate sets absolute protein abundance regulation across cell types

One property that has been suggested to influence the mode of gene expression regulation is growth rate^12,19,20^. Some reports have emphasized the compensation of protein dilution by changes in synthesis rates^19,20^ while others have emphasized its impact on altering protein abundance^12^. No estimates exist for primary cell types directly from mammalian tissues. To investigate this influence, we first estimated cell growth rate based on the incorporation of the metabolic label into histone proteins^32^. Here, we explicitly computed the effective doubling time of each cell type by counting the number of cell divisions that occurred since the feeding started, see methods Fig. 2a. Cell types displayed a variety of growth rates ranging from 9.9 day doubling time to no recorded division events within the 10 day pulse.

These growth differences are important since the growth related dilution contributes to protein clearance. At faster growth rates, the rate of dilution assumes a larger fraction of protein clearance, reducing the protein to protein variability in clearance rates^12,19^, Fig. 2b,c. As a result, the dynamic range of clearance rates is reduced (the smallest rate is the rate of cell growth), thus directly affecting the scaling of clearance rates between cell types, as visualized via the shallow slope when comparing clearance rates between cell types with growing at different rates, Fig. 2b. The dependence of this slope scaling with growth rate is strongly confirmed by the fact that the slope between degradation rates for these cell types is close to 1, Extended Data Fig. 7a.

Independent from the slope scaling, the data reveal significant variation (scatter) of the clearance rates of basal cells and chondrocytes, with a correlation of only 0.50, despite higher reliability of the clearance rate estimates (0.7 - 0.75). This trend is further observed between most cell types, Extended Data Fig. 3i. This variation reflects cell-type specific differences in growth rates, as indeed confirmed by the fact that the correlation between the corresponding protein degradation rates is the same.

**Figure 2.**
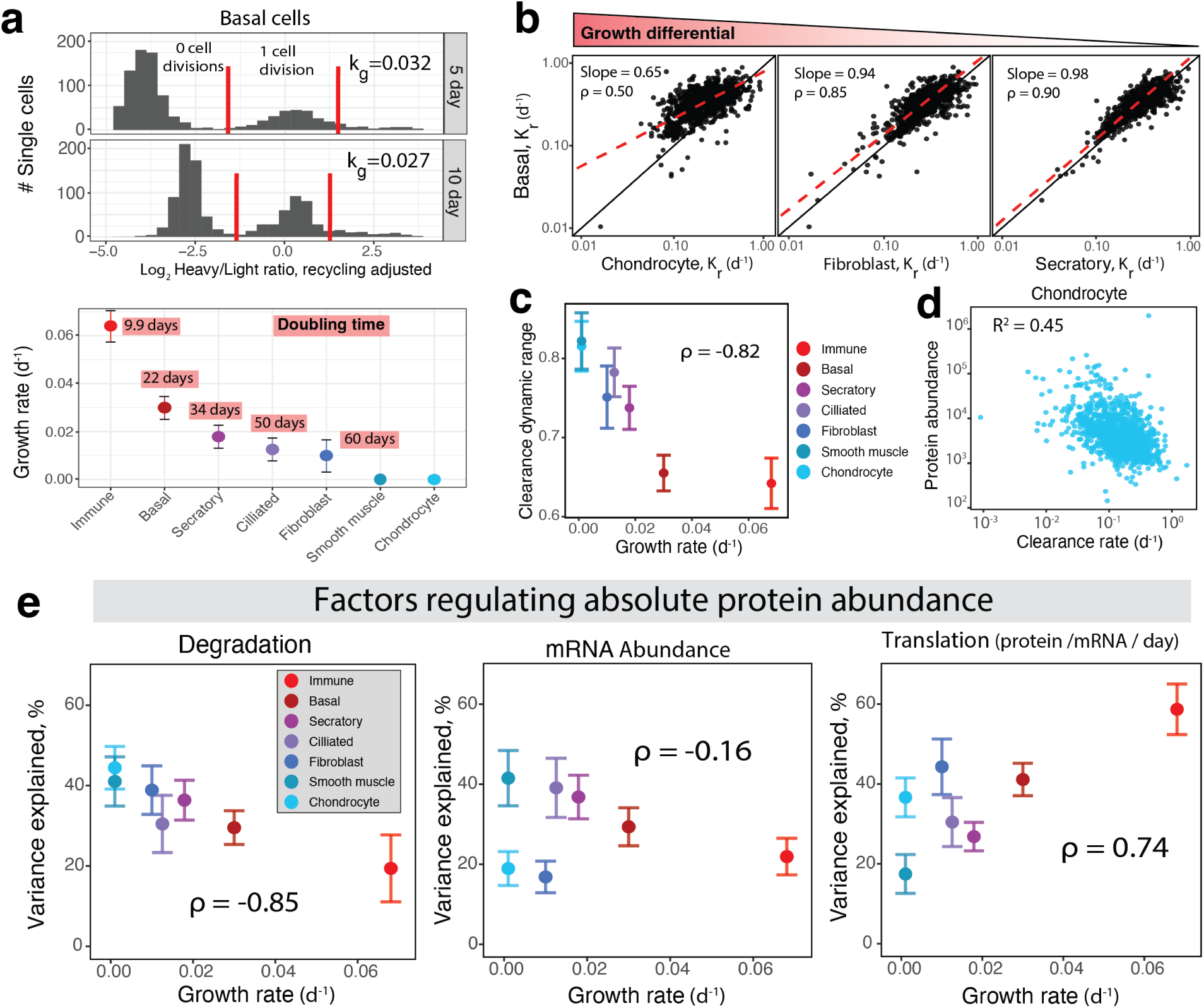
The regulatory contributions of translation and clearance to absolute protein abundance scales linearly with growth rate. **a**, Plotting the recycling adjusted heavy to light ratios of histone peptides reveals multimodal distributions, with modes corresponding to the number of times cells have divided and partitioned histones. These distributions were used to infer cell doubling time, which is proportional to growth rate, *ln*(2)*/k_g_*. **b**, Comparisons of clearance rates between different cell types, ordered by growth-rate differences between the cell types. For cell types with more similar growth rates, the slope of the clearance rates between the cell types converges to 1. **c**, The variability in degradation rates across proteins is estimated by the standard deviation of the log transformed clearance rates and plotted against growth rate, *r* = −0.82. **d**, For the most slowly growing chondrocyte cells (with no recorded cell divisions), protein clearance explains 0.45 % of protein abundance variation. **e**, Growth rate strongly predicts the influence of degradation and translation on absolute protein abundance. The influence of translation is proportional to the growth rate while the influence of clearance is inversely proportional.

In addition to cell-type specific degradation, clearance rates have a wider dynamic range at low growth (Fig. 2b), and thus the potential for stronger impact on protein abundance. To evaluate this potential across all cell types, we compare the dynamic range of the clearance rates variation across proteins to the corresponding cell growth rate. We found a strong inverse relationship with growth rate, *ρ* = −0.82, Fig. 2c, consistent with the expectation. To more directly quantify if the potential is realized, we examined the joint distribution of clearance rates and protein abundance in chondrocytes. It shows a protein clearance explains roughly 45 % of the variation in protein abundance estimated by square of the Pearson correlation, Fig. 2d. Conversely, only 29% of variance is explained for faster growing basal cells, Extended Data Fig. 7b. Overall, the extent of abundance variation explained by clearance attenuates as growth rate increases, following a clear linear dependence Fig. 2e. This result extends our previous observation made across tissues to cell types within a tissue^12^.

Accurately estimating this influence required accounting for reliability of the measurements. To do so, we applied the Spearman correction to the fraction of variance explained based on reliability estimates derived in Extended Data Fig. 6e, see methods. We also applied this method to estimating the fraction of variance explained by mRNA abundance, Fig. 2d. While both mRNA abundance and clearance are estimated relatively independently of protein abundance, translation shares peptide ionization bias with protein abundance estimates. Thus, estimates for fraction of variation explained by translation were taken to be the remaining variance. These estimates are in agreement with another estimate derived from the square of the Pearson correlation, Extended Data Fig. 7d. However, it should be noted that any estimation of translational regulation depends on the accuracy of our data reliability corrections. To assess the robustness of the relationship between translational regulation and growth rate, we also examined estimates made without correction, Extended Data Fig. 7e. We then performed a sensitivity analysis by adding random gaussian noise to reliability estimates made for each cell type, Extended Data Fig. 7f, see methods. Our sensitivity analysis revealed an average correlation between growth rate and translational contribution to protein variation of 0.61. This correlation is slightly lower but qualitatively recapitulates the scaling of translational regulation with cell growth rate. The contribution of mRNA abundance did not correlate strongly to growth rate. In contrast, translational regulation increased in direct proportion to growth rate, Fig. 2d.

### Growth rate sets absolute protein abundance regulation within basal cells

Having established that cell growth rate shapes the regulation of absolute protein abundance across cell types, we asked whether the same principles extend to continuous variation within a single cell type. Basal cells were a natural choice, as previous work has reported a proliferative subpopulation of hillock basal cells marked by high Krt13 protein abundance^18^. Consistent with this, Krt13 and its heteropolymer-forming partner Krt4 formed one of the most strongly co-varying protein pairs across basal cells, *r* = 0.59, Extended Data Fig. S1a. We hypothesized that Krt13 abundance might mark an axis along which cellular growth rate increases, allowing us to compare growth rate across the protein and mRNA modalities with greater resolution and confidence.

To test this, we first aligned basal-cell protein and mRNA measurements along a common axis by projecting mRNA cells into a principal component basis learned from 1,345 protein cells, see methods, Fig. 3a. The first common principal component (PC1) captured a shared axis along which the Krt13/4 signature increased in both modalities, *r* = 0.96 at the protein level and *r* = 0.98 at the mRNA level, Fig. 3a. This axis coincided with cell division: the fraction of cells that had divided over the labeling period rose from roughly 20% to over 60% along PC1, *r* = 0.94, Fig. 3b, supporting our hypothesis that Krt13 abundance can serve as a proxy for cell growth rate^18^.

We next asked whether the regulatory shifts observed across cell types recurred along this within-basal-cell axis. The correlation between protein clearance and abundance decreased across PC1 bins, *r* = 0.9, Fig. 3c, consistent with faster-growing cells relying less on clearance regulation and in direct alignment with our observations at the cell-type level. In parallel, the average protein-mRNA correlation increased along PC1, *r* = 0.64, Fig. 3d, indicating stronger transcriptional coupling in the more proliferative state. This positive scaling between transcriptional coupling and growth rate is stronger than the corresponding relationship across cell types, indicating that growth-associated transcriptional regulation is more prominent along the within-basal proliferative axis than across cell-type boundaries.

While Krt13 protein and mRNA co-varied along the common axis, indicating that its abundance is set primarily by transcription, we next asked how the broader molecular machinery driving proliferation was regulated along the same axis. Sustaining rapid growth requires cells to synthesize substantial amounts of new protein, and consistent with this, total protein synthesis was tightly coupled to cell division along the axis, *r* = 0.96, Supplemental Fig. S1b, supporting the growth-rate interpretation of this axis. We then examined how the machinery of protein synthesis and its energetic support scaled with total synthesis.

Ribosomal proteins, the proteins directly involved in translation, rose sharply with total protein synthesis, *r* = 0.98, while ribosomal mRNA levels moved in the opposite direction, slightly declining, *r* = −0.92, Fig. 3e. Because protein synthesis is one of the most energetically expensive cellular processes, we also asked how mitochondrial proteins, which supply ATP via oxidative phosphorylation, scaled along the same axis. Mitochondrial proteins showed the opposite trend, decreasing along the synthesis axis at both the protein and mRNA level, though more strongly at the protein level, protein *r* = −0.9 vs mRNA *r* = −0.7. This apparent shift away from oxidative energy production despite rising translational demand mirrors the classic Warburg switch, in which proliferating cells favor glycolysis over oxidative phosphorylation for both ATP and the biosynthetic precursors needed to build biomass. We therefore asked whether the glycolytic enzymes along this axis followed the same pattern.

**Figure 3.**
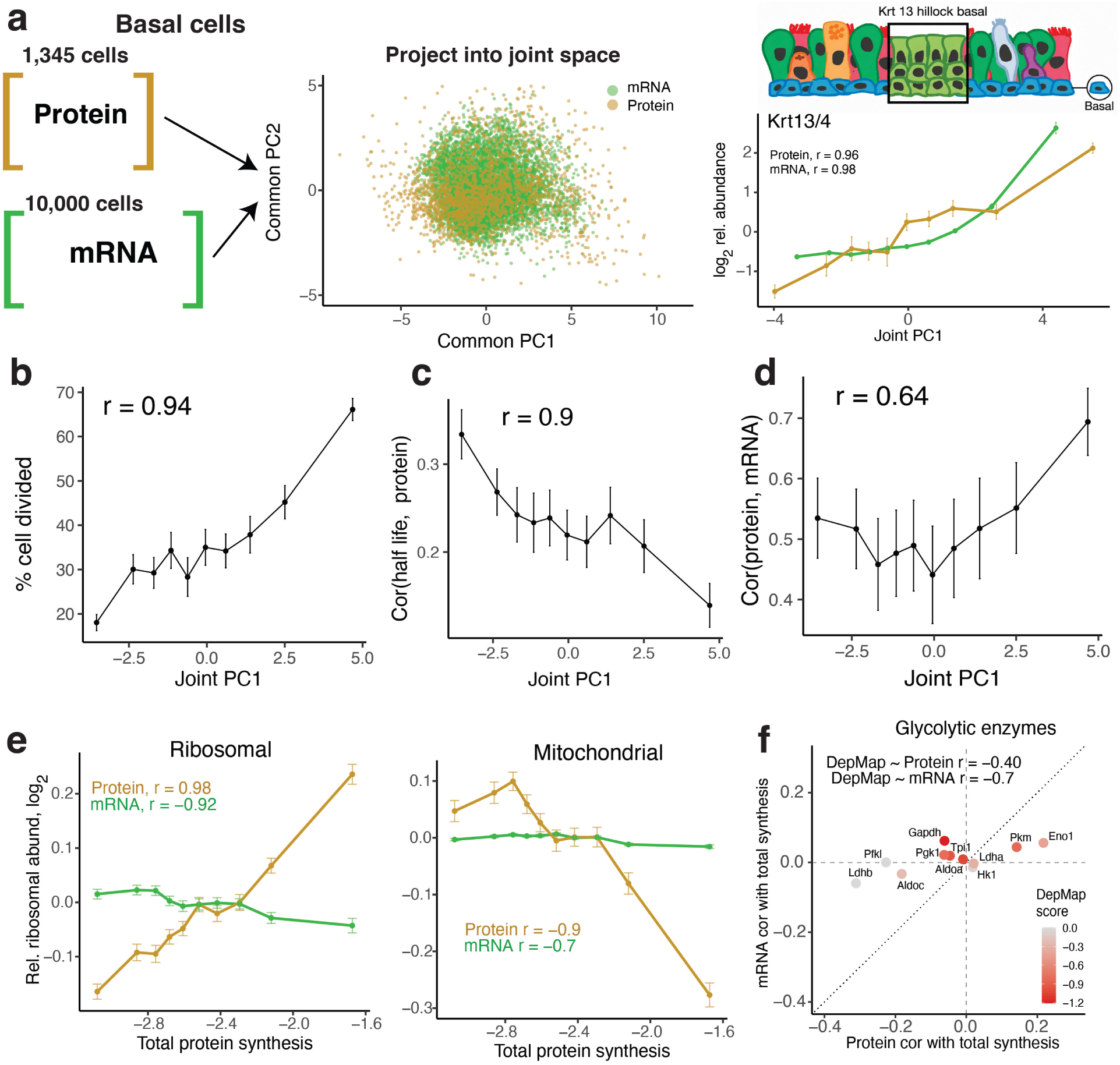
Growth rate gradient within basal cells affirms the protein regulation principles. **a**, To compare protein and mRNA levels within basal cells, we projected the data into a common space using common principal component analysis. This space captured one known axis of intra-basal variability with the increase in Krt13 expression corresponding to hillock like basal cells^18,29^. **b**, Percentage of cells that have undergone cell division within 10 equally sized bins spanning the first common principal component. **c**, The correlation between average absolute protein levels and clearance rates for cells within each bin plotted across the first principal component. **d**, The correlation between average absolute protein levels and mRNA levels for cells within each bin plotted across the first principal component. **e**, Relative ribosomal protein abundance strongly increase with total protein synthesis while ribosomal mRNA abundance shows a slight decrease. Similarly, protein abundances for mitochondrial genes significantly decrease compared to mRNA abundances for the same set of genes. **f**, The correlation of glycolytic enzymes to total protein synthesis varies across different gene products. Correlation to total protein synthesis is inversely correlated to DepMap essentiality^45^ at the level of both protein and mRNAs.

Glycolytic enzymes similarly exhibited a Warburg-like isozyme switch. Cancer-associated glycolytic isoforms (Pkm, Eno1) increased with total protein synthesis while oxidative isoforms (Ldhb, Pfkl) decreased, Fig. 3f. Coloring these enzymes by their DepMap CRISPR proliferation dependency scores^45,46^ revealed strong agreement: enzymes most essential for cancer cell proliferation (Pkm, Eno1, Ldha) were upregulated in high-synthesis basal cells, while dispensable isoforms (Ldhb, Pfkl) were downregulated. To test whether this correspondence generalizes, we computed per-gene protein and mRNA bin-trend correlations with total synthesis across all measured genes and correlated them against DepMap dependency, Supplemental Fig. S1c. We observed a genome-wide agreement stronger at the protein level than the mRNA level, protein *r* = −0.25 vs mRNA *r* = −0.1, indicating that DepMap-associated proliferation essentiality is preferentially recovered by protein-level measurements.

### Regulation of relative protein abundances across cell types

Having analyzed how different regulatory modes influence absolute abundance variation across protein , we next used our measurements to characterize how individual proteins are regulated across cell types. Similarly, we aim to quantify the impact of all regulatory modes. The simplest possibility is that the relative abundance of a protein is regulated primarily by a single mechanism. This is illustrated with Sult1d1, which primarily co-vary with mRNA abundance, *ρ* = 0.98, Fig. 4a. Other proteins such has Sec12I2 primary co-vary with translation, *ρ* = 0.98. To estimate measurement reliability, we leverage the quantification of distinct peptide sequences for a given protein along with sample preparation replicates within a Bayesian framework. This allowed us to reflect higher uncertainty in the translation quantities, which share uncertainty from the mRNA abundance, protein abundance, and protein clearance measurements, see methods. Our uncertainty estimates control for factors such as sample preparation variability and potential MS interferences.

To explore the full range of regulation for all quantified proteins, we correlated protein abundance to mRNA abundance, translation rate, and clearance half-life, Fig. 4b and Fig. 8a. The correlations for all regulatory modes were broadly spread and positively biased, indicating that proteins are regulated by different modes to different degrees. The average correlation was the highest for mRNA abundance, suggesting it significantly influences the abundance of many proteins. For some protein groups, such as actin filaments, RNA abundance appears the dominant regulatory factor. For other protein groups, translation and protein clearance appear the dominant regulatory factors, Fig. 4b. Specifically, different pathways controlling metabolic functions, translation, localization, and degradation of proteins were primarily controlled by translation and clearance. We further extended this analysis to important functional groups of proteins such as chromatin binding proteins, mRNA binding proteins, and E3 ligase, Fig. 4c. Clearance played a strong role in regulating relative levels of mitochondria and the proteosome, while translation had a prominent regulatory role for E3 ligases. The relative influence of translation and clearance has only previously been estimated across a single cell type cultured in vitro and responding to stimulus^10^. Our results suggest stronger overall regulatory impact when extending measurements to the new context of how diverse cell types from a primary tissue regulate their proteomes.

The univariate analysis in Fig. 4, suggests that most proteins are regulated by multiple mechanisms, and we further investigated this with bivariate analysis, Extended Data Fig. 8. We observed that as the correlations between proteins and mRNA abundance decreases, the correlation between protein and translation or clearance rates increase, Extended Data Fig. 8a. Interestingly, proteins with the strongest correlation to mRNA abundance had a strong inverse correlation to clearance half-life. This suggests cell type specific proteins with fast turnover require even tighter transcriptional control. However, these mechanisms may not only act independently, but synergistically. Indeed, proteins such as St13 are co-regulated by both transcription and translation, Extended Data Fig. 8b. This can also be observed for groups of proteins, such as certain keratin filaments synergistically regulated by both transcription and translation in basal cells Extended Data Fig. 8d. The contributions of all modes to the regulation of proteins abundance is listed in supplementary table 1, both for individual proteins and GO terms. Collectively, our results indicate that the dominant mode of regulation across cell types varies across proteins.

The primary mode of regulation also varies across cell types. For example, while protein clearance explains 38% of variation in relative protein abundance between basal cells and chondrocytes, it explains little variation for the same set of proteins between different cell types, such as basal and immune cells, Fig. 4d. Generally, we see clearance explain the most significant amount of variance between cell types with slower growth rates such as chondrocytes and smooth muscle cells. This is because cell-type specific differences in degradation exert greater effect on protein clearance when clearance rates are less influenced by protein dilution.

**Figure 4.**
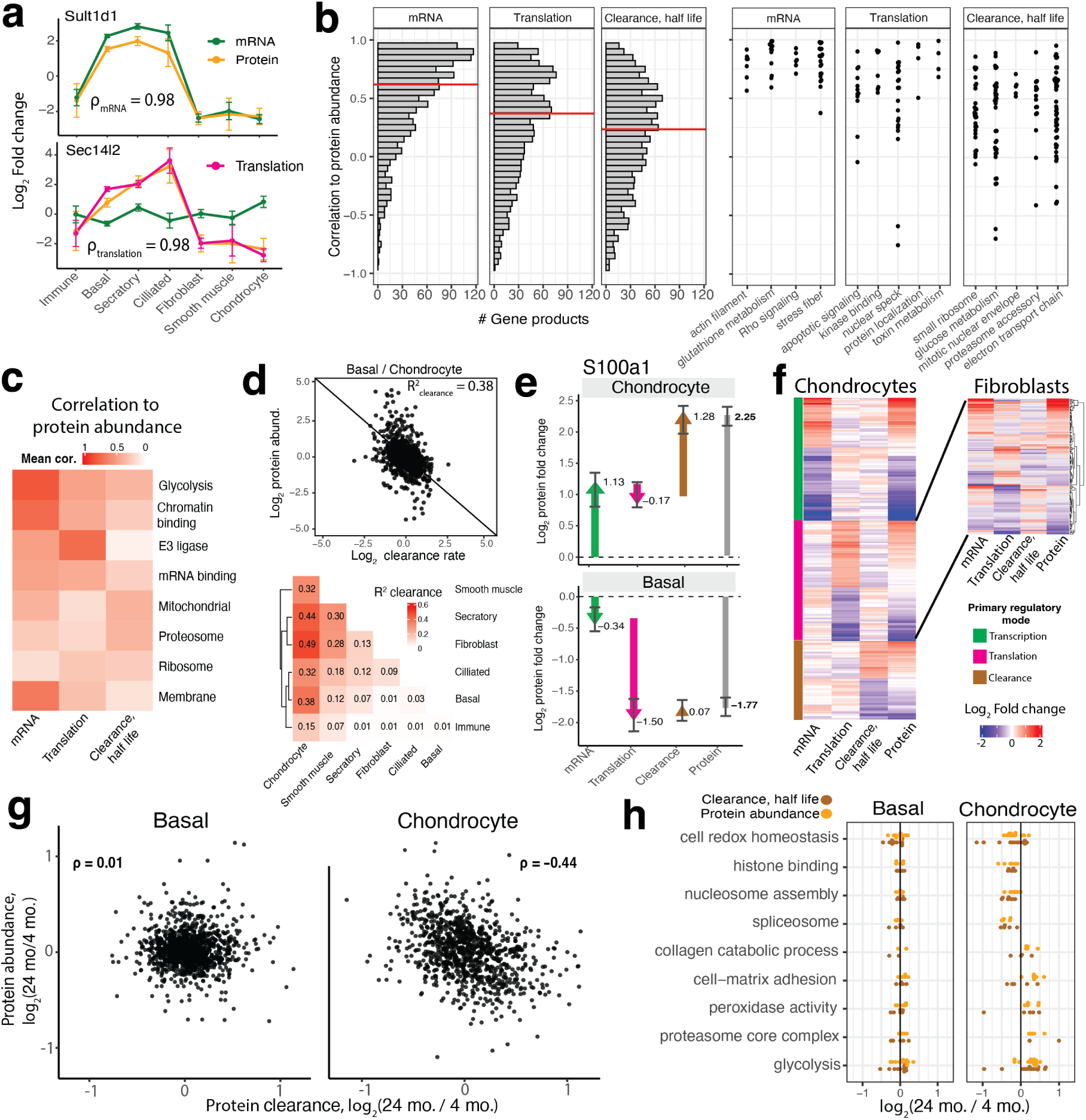
Regulation of relative protein abundance across cell types. **a**, Examples of proteins regulated by a primary mechanism across all cell types, mRNA abundance for Sutl1d1 and translation control for Sec12I2. **b**, Correlations of cell-type average protein abundance to mRNA abundance, translation rate, and protein clearance half lives for all proteins. Example proteins predominantly controlled by mRNA abundance, translation, or clearance are plotted, displaying correlations for individual proteins grouped by GO terms. **c**, Average correlations for each mode of regulation to proteins from functional groups. **d**, Clearance contributes significantly to protein fold changes between basal cells and chondrocytes, but less for other cell-type pairs growing faster, as shown in the heatmap. **e**, Examining protein S100a1 demonstrates how regulation methods can be cell type specific, with mRNA abundance and clearance explaining high abundance in chondrocytes, but low translation explaining low abundance in basal cells. **f**, Heatmaps are displayed for proteins organized by the primarily regulatory mode, mRNA abundance, translation, or clearance. For proteins primarily regulated by translation in chondrocytes, there is substantial diversity in regulation patterns in fibroblasts. **g**, Comparison of protein abundance and clearance fold changes for old and young mice. A replicate comparison is shown in Extended Data Fig. 9. **h**, Protein abundance and protein clearance fold changes between old and young mice for functional groups of proteins.

Next, we increase our resolution and consider the variation in the regulatory mode both across cell types and proteins. As an example, S100a1, is up-regulated in chondrocytes because of increased RNA abundance and decreased clearance while in basal cells, the reduced abundance is primarily achieved by translational suppression, Fig. 4e. To visualize such trends, we plotted the subsets of proteins whose relative abundance in chondrocytes is primarily regulated by a single mode of regulation, Fig. 4f, see methods. To explore whether regulation is protein-specific and similar across cell types, we focused on the subset of proteins primarily regulated by translation in chondrocytes. Visualizing their regulatory modes fibroblasts shows a mix of regulatory strategies, including significant control by transcription and clearance, Fig. 4f. These results generalize the observations from S100a1 and indicate that the mode of regulation is both protein and cell-type specific.

To visualize this result more globally and representatively, we systematically identified functional groups of proteins whose regulation shows strong enrichment for transcription, translation, or clearance, Extended Data Fig. 8d. Some terms such as cytoplasmic translation were regulated by different modes across cell types such as transcription in basal cells and translation in secretory and immune cells. Many structural process were regulated by clearance across cell types such as microtubule polymerization in ciliated cells, lamellipodium organization in smooth muscle cells and intermediate filament organization in immune cells.

Next, we investigated protein abundance regulation across age groups. To assess the extent of reproducible differences in regulation for each cell type, we first compared protein fold changes across replicate mice at ages 4 and 24 month. We found that only chondrocytes had reproducible protein changes with age, Extended Data Fig. 9a. Similarly, only chondrocytes exhibited significant and reproducible differences in protein clearance, Extended Data Fig. 9b. The strong dependence of clearance rate on age for chondrocytes may reflect that these cell types have the lowest recorded rate of cell division. One potential implication of low cell division is that aberrant or misfolded proteins have greater potential to accumulate as they are not diluted by cell division contributing to more significant proteostasis stress^47^. However, the limited number of aged mice should be noted and future experiments would be needed to validate this finding. To understand how protein abundances differences are regulated across chondrocytes from old and young mice, we compared fold changes of protein abundance and clearance rates, Fig. 4g and Extended Data Fig. 9. The agreement between abundance and clearance fold changes, *ρ* = −0.44, suggests substantial contribution of age associated changes in protein degradation rates. No other cell types showed agreement between clearance and protein fold changes across age as exemplified by basal cells, Fig. 4g.

Despite the loss of old mice, the data indicate significant enrichment of functional groups of proteins with relation to age. Protein set enrichment analysis across chondrocytes from young and old mice found 123 functional groups significant at 5% FDR, supplemental table 2. This is contrasted with just 1 significant functional group for basal cells. Among these sets of proteins were histone binding and nucleosome assembly up-regulated in young mice, Fig. 4h. We also observed that proteins participating in redox homeostasis were up-regulated in young mice while proteins participating in peroxidase activity and glycolysis were up-regulated in old mice.

### Regulation of mRNA and protein covariation within cell type

Although grouping cells by type is convenient and explains much of the protein abundance variability, cells vary functionally even within a cell type. These functional variations are mediated in part by regulated protein abundance changes. Indeed, when examining the most numerous basal cell population, we found that some proteins such as Cavin1 varied at least 30 fold, Fig. 5a. Further, the abundance of Cavin1 varied uni-modally across single cells, suggesting that further sub-clustering would incompletely explain its variation. Interestingly, mRNA levels for Cavin1 varied about 4 fold less. This difference in dynamic range for the protein and RNA products of a gene generalize to all analyzed proteins: the protein abundances vary to a greater degree across single cells than mRNA for over 90 % of quantified genes products, Extended Data Fig. 10a. To better understand the implications of this variation and distinguish between biological and technical origins, we sought to analyze covariation patterns to identify potential functions associated.

Further, as we did not measure mRNA and protein abundance in the same single cells, comparing single-cell covariation patterns at the protein and mRNA level enabled us to determine the extent to which protein level covariation is regulated transcriptionally.

The large variation of Cavin1 within basal cells is likely reflecting the variable abundance of caveolae structures, as supported by the strong correlation (*ρ* = 0.81) to the other subunit, Cav1, Fig. 5a. At the mRNA level, we observed a weaker correlation between Cavin1 and Cav1 of *ρ* = 0.34. One possible reason for this correlation difference might be the uneven partitioning of cellular content during cell division imprinting more strongly on proteins due to their longer half lives. We excluded this possibility by confirming that cell division history did not explain Cavin1 or Cav1 variation, Extended Data Fig. 10b. Furthermore, significant protein-protein correlations and variation of protein abundance within a cell type are observed in non-proliferative cell types, such as chondrocytes, indicating that these phenomena cannot be attributed to cell division. Extending this analysis to all protein pairs indicates an overall similarity (Pearson correlation) between pairwise correlations of RNA and proteins, Ψ = 0.31. Beyond the similarity of covariation at the mRNA and protein level, this result emphasizes significantly stronger correlations among protein pairs. For example, over 5,000 pairwise protein correlations exceed a correlation of 0.3, compared to just 340 at the mRNA level. Again, we observed that cell division history did not explain the magnitude or nature of protein-protein correlations, Extended Data Fig. 10c.

To determine whether these high correlations reflected cellular functions, we compared groups of related gene products. Proteins annotated to interact in either the CORUM or STRING database showed significantly higher correlation at the protein level as comapred to mRNA, Fig. 5c and Extended Data Fig. 10e, see methods. These results suggest that protein correlations capture functional covariation and motivated us to examine differences in clustered correlation modules, however, it should be noted generally that strong covariation does not provide direct evidence of interaction. We applied hierarchical clustering on a subset of proteins and mRNAs, see methods. First, this subset consisted of proteins and mRNAs for which the proteins had at least one pairwise correlation above 0.3. This resulted in a group of roughly 400 unique proteins. We then plotted the correlation patterns against each other as the lower and upper triangle of a correlation heatmap^24^, Fig. 5d and Extended Data Fig. 10f. We observed several distinct correlation modules primarily present at the protein level, each found to be enriched for different functional processes, FDR *<* 0.01. For example, a module of proteins enriched for mesenchymal transition had an average correlation of 0.43 at the protein level and 0.02 at the mRNA level. Applying the same procedure, but on a subset of gene products where the mRNAs contained at least one pairwise correlation above 0.2, we found fewer confident clusters and many had no clear functional enrichment, Extended Data Fig. 10f.

We then set out to understand why certain covariation patterns would be present primarily at the protein level. One potential reason is that active regulation of protein translation or clearance contribute to the functional protein covariation not observed at the level of transcripts. As we measured protein clearance rate with single-cell resolution, we investigated its contribution. To gain intuition, we first inspected the chromatin bound proteins Hmga1 and Hmgn2 whose correlation at the protein level is 0.56, while only 0.01 at the level of transcripts Fig. 5d. This difference may stem from the regulation of Hmga1 protein abundance by clearance, reflected in a correlation of *ρ* = −0.48 between Hmga1 clearance rates and abundance across cells, Fig. 5d. To explore if this example is representative and clearance regulation contributes globally, we plotted the difference in correlations between mRNA and protein abundance, versus the extent to which each protein was regulated by clearance (the average of the correlations of clearance and abundance for each protein across single cells). We observed that pairs of gene products more highly correlated at the protein level are significantly more likely to be controlled by protein clearance, Fig. 5f. Our predictive power improved from *r* = 0.46 to *r* = 0.55 when we also considered the absolute half life of each protein in a linear model, Extended Data Fig. 10h. This may be due to the fact that for longer lived proteins, the comparatively stochastic expression of short lived mRNA poorly reflect protein abundance. To ensure these trend were not induced by systematic differences between estimates of clearance from mice pulsed for 5 or 10 days, computed this relationship across cells that have only been pulsed for 5 days and observed a similar trend, Extended Data Fig. 10g.

We then expanded our analysis to test whether these trends generalize to other cell types and depend on factors such as growth rate. We constrained our analysis to populations with at least 500 single cells to ensure a sufficient number of data points when computing protein-protein correlations. To estimate the contribution of clearance to protein-protein correlations, we compared the pairwise correlations between protein abundance and clearance and quantified the similarity by the correlation Ψ, analogous to Fig. 5b; for chondrocytes, clearance Ψ = 0.24, suggesting that protein clearance contributes significantly to shaping protein-protein correlations, Fig. 5g. Analogously, we computed for other cell types clearance Ψ and RNA Ψ (correlation between RNA-RNA and protein-protein correlations), and plotted them as a function of the growth rate, Fig. 5g. The results indicate strong linear dependencies: For more slowly growing cell types such as chondrocytes, clearance covariation more strongly reflected protein abundance covariation. This trend was reversed when comparing protein and mRNA covariation patterns to cell growth rate. These results were consistent with our previous observations about more slowly growing cells relying more strongly on regulation by protein degradation. However, the extent of regulation by transcription within cell type depends more on growth rate than observed at the absolute abundance level.

**Figure 5.**
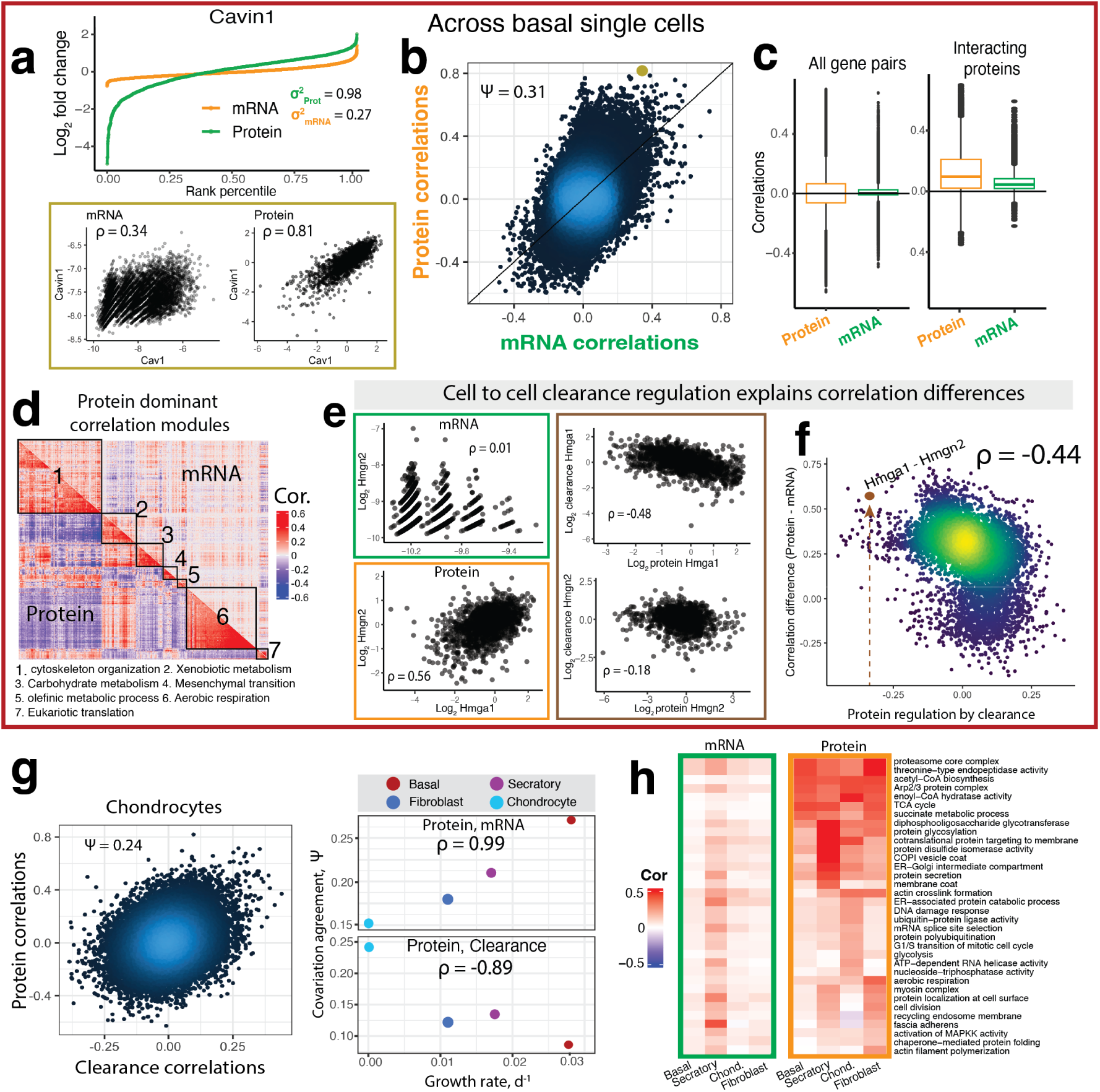
Protein covariation within cell type is only partially reflected in mRNA covariation and significantly shaped by protein clearance regulation. **a**, The dynamic range of measured protein concentrations for Cavin1 is about 4 fold larger than at the mRNA level. Pairwise scatter plots of relative fold changes across single basal cells for Cav1 and Cavin1 transcripts and proteins. **b**, A scatter plot of all pairwise protein-protein versus corresponding mRNA-mRNA correlations reveals a correlationΨ = 0.31. **c**, Pairwise correlations for all proteins and transcripts are centered around 0, but correlations for interacting proteins are positive and larger than the corresponding mRNA correlations on average 0.12. **d**, Protein correlation modules detected via hierarchical clustering are plotted with correlations for the corresponding transcripts. **e**, Proteins Hmga1 and Hmgn2 are correlated but their mRNAs are not. This discrepancy is partly explained by the clearance regulation of these proteins by clearance reflected in the correlations between abundance and clearance rates. **f**, The extent of protein clearance regulation for protein pairs can explain the difference between their correlation and the correlations among corresponding mRNA pairs, *ρ* = −0.44. **g**, Pairwise protein clearance rate correlations are similar to the corresponding protein abundance correlations, Ψ = 0.24. This agreement Ψ depends linearly on the cell growth rate. The agreement between pairwise protein and mRNA correlations exhibits the inverse linear dependence. **h**, GO terms whose members covary significantly, FDR of 1 % for proteins and mRNA. Proteins show considerable differences across cell types while differences in covariation of mRNA levels are less present.

Having established differences in correlation patterns for proteins and RNAs across single cell, we next examined the differences in covariation patterns within functional groups of proteins and mRNAs. For each cell type, we computed the protein-protein and mRNA-mRNA correlations within a functional group and compared them to the corresponding distribution of all correlations within each modality. We observed 315 GO terms with significantly (at 1% FDR) higher protein correlations in at least one cell type. In contrast, only 185 GO terms reached significance at the mRNA level (supplemental table 3, Fig. 5h), despite the larger number of single cells. These differences suggest that protein abundance variation is better at capturing functional state variation across individual cells. These functional states show significant degrees of cell-type specificity, as demonstrated by the series of different functional groups most strongly co-regulated within individual cell types, Fig. 5h. Notably, terms such as ER-Golgi intermediate compartment and other terms involved in protein secretion were significantly correlated in the secretory cell population, while proteins participating in glycolysis strongly co-varied in the chondrocyte cells inhabiting low oxygen environments,^48^.

### Cell type specific single cell protein covariation patterns

Having observed strong covariation patterns at the protein level, we sought to further leverage this signal to infer differences in the coordination of proteins functions across cell types. We originally observed large dynamic range spanning 50 fold for the concentration of intermediate filaments (IF) in both basal cells (Keratin 5) and fibroblasts (Vimentin). Curious as to why these cells would have such varied concentrations, we explored which proteins were correlated to high IF concentration by computing the correlations of all proteins to Krt5 and Vim in both fibroblast and basal cells, i.e., correlation vectors. Interestingly, for both Krt5 and Vim, correlation vectors showed no similarity between basal cells and fibroblasts, Fig. 6a. However, the correlation vectors of the primary IF in each cell type (Krt5 in basal cells and Vim in fibroblasts) are strongly correlated, *ρ* = 0.80. This correlation pattern illustrates how correlations can reveal functional changes.

To understand the functional variability associated with high or low IF concentration, we next inspected the subset of proteins with high positive and negative correlations to the primary IFs, Fig. 6b. We observed a gradient of protein abundance present in both cell types. It reflects a shift from cells with high concentration of cytoskeletal proteins (suggesting high mechanical force) to cells with high concentration of proteins participating in cellular redox response to stress. These results suggest a common phenotypic axis of variation, with unique proteins participating from each cell type based on the similarity of their primary functional role contingent on cellular context. We sought to generalize this analysis to identify additional proteins that potentially change their functional co-ordination across cell types. The first step in our procedure was to compute vectors of pairwise correlations for the i*th* protein, denoted as **b***_i_* in basal cells and **f***_i_* for fibroblasts. We then quantified the similarity between these correlation vectors by their correlations, ⟨**b***_i_,* **f***_i_*⟩ as previously described^49,50^, Fig. 6c, supplemental table 4. The distribution of correlations was on average positive, mean *ρ* = 0.21, suggesting shared covariation patterns between cell types. However, many proteins showed minimal correlation vector similarity across cell types. We then partitioned the proteins into two sets of highly similar and dissimilar proteins with a threshold for high (*ρ >* 0.50) and low (*ρ <* 0.10) similarity. High similarity proteins from set 1 constituted an axis of common covariation across cell types, including groups of proteins participating in cell metabolism, cytoskeleton, and translation, Fig. 6c.

For the proteins in set 2 with low agreement in their pattern of covariation across single cells, Fig. 6c, we plotted the correlations between set 1 and set 2 proteins within each cell type. By clustering correlations between set 1 and set 2 proteins for each cell type, we identified opposing covariation patterns, Fig. 6d. In basal cells, set 2 proteins associated with intermediate filament and fatty acid metabolism (purple cluster) positively correlated with set 1 proteins related to the TCA cycle and cellular stress, while proteins associated with glycoprotein metabolism and cell adhesion (green cluster) correlated positively with set 1 proteins involved in translation and glycolysis. These trends were reversed in fibroblasts, Fig. 6d. These opposite patterns of covariation raise the possibility that, for example, basal cells rely on TCA and oxidative phosphorylation to generate energy for intermediate filament generation while fibroblasts rely on glycolysis. However, determining the nature of this dependence requires further validation such as genetic perturbations or joint analysis of metabolites and proteins in single cells.

**Figure 6.**
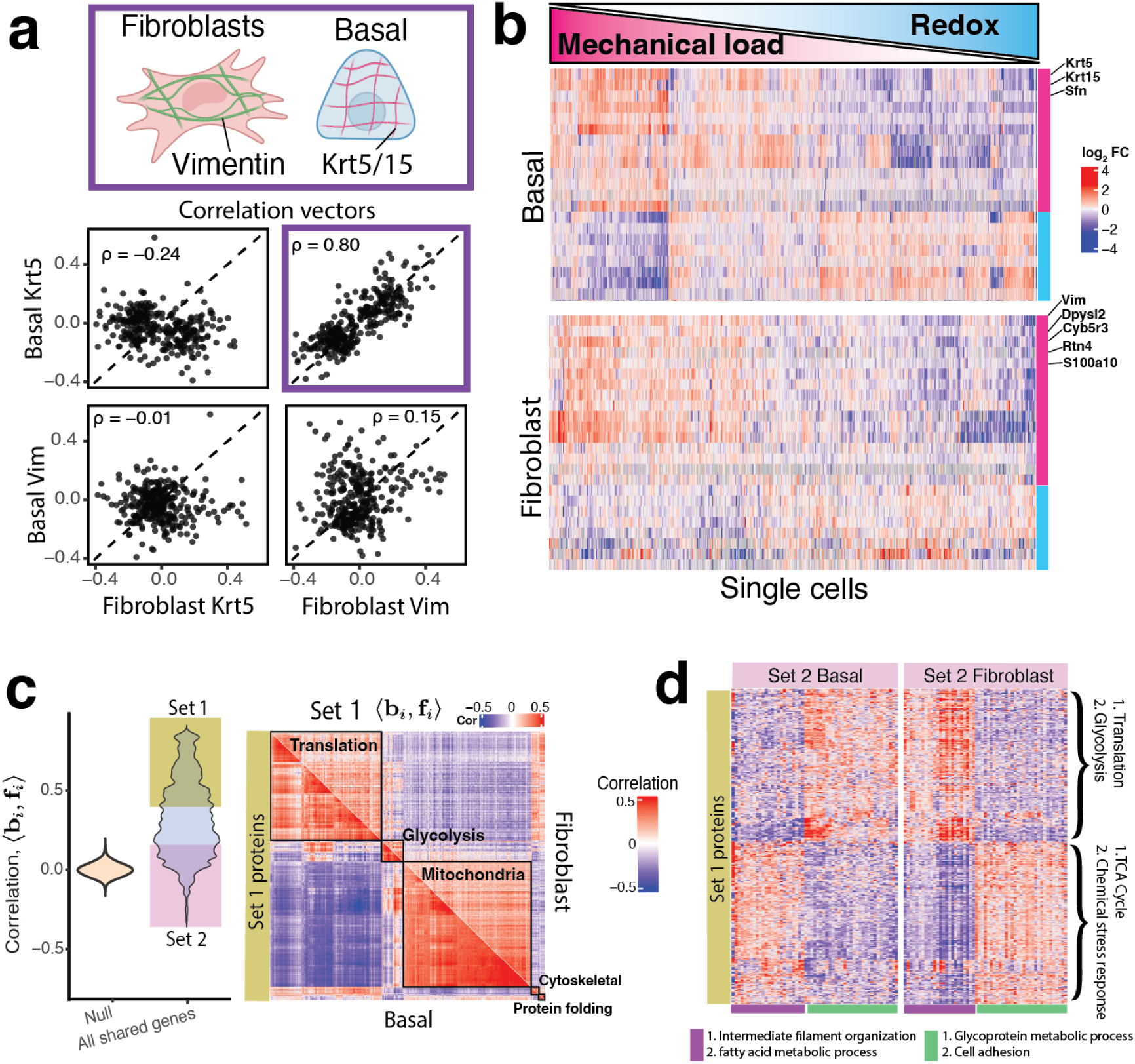
Cell-type specific covariation identifies functional exchanges between proteins. **a**, Vimentin is the primary intermediate filament (IF) in fibroblasts while Krt5/15 are the primary IFs in basal cells. The correlation vectors (to all other proteins) of each IF are cell-type specific and similar between basal and fibroblasts cells. **b**, Protein abundance across single cells shows gradients of cells defined by the co-expression of either cytoskeletal proteins (suggesting cells are under higher mechanical strain) or redox proteins (suggesting high oxidative stress). **c**, The correlations between correlation vectors for all proteins across basal and fibroblast cells are broadly distributed with many proteins showing high covariation (set 1) while fewer proteins showing low covariation agreement (set 2). A clustergram of protein proteins from set 1 defines functional clusters. **d**, Clustergrams of pairwise correlations between proteins from set 1 and set 2 identify clusters (marked by purple and green) with changing correlations between basal and fibroblast cells.

To understand how these differences in protein-protein correlations are achieved, we examined the potential for regulation by transcription and protein clearance. Applying the same correlation vector analysis to the corresponding mRNA and clearance rate data resulted in much weaker correlation structures, Supplemental Fig. S3a. To reduce the potential contribution of measurement noise, we compared the co-variances of the averaged mRNA and clearance rates across proteins from the two clusters of set 1 and set 2 proteins, Supplemental Fig. S3b. This revealed correlation structures in the mRNA abundance and protein clearance similar to the structure of protein-protein correlations This similarity suggests that both RNA abundance and protein clearance contribute to the protein clusters in Fig. 6d.

## Discussion

Our data and analysis allowed us to extend the pioneering studies of gene expression regulation in vitro^9,10^ to the single cells comprising a complex tissue. This expansion revealed new regulatory trends. Our results confirm substantial translational regulation at fast growth rates, consistent with previous observations in rapidly proliferating cells^9^. However, by examining cell types spanning a broad range of growth rates, we uncover a progressive shift in the predominant mode of protein abundance regulation from translation at fast growth to protein clearance at slow growth, with clearance eclipsing translation in terminally differentiated cells.

We observe remarkably simple linear scaling of the contributions of translation and protein clearance to protein abundance variation. This simplicity of the linear scaling is reminiscent of scaling laws that have been observed as a function of cell growth rate in bacteria^51^ and yeast^52^. Crucially, we can mechanistically explain the increasing role of protein clearance with the decreasing dilution rate at slower growth rates, if (and only if) this dilution effect is not compensated by adjustment of synthesis rates as previously posited^12,19,20^. In contrast, the scaling of the translation contribution remains phenomenological but consistent with previous estimates from rapidly growing cells in vitro^9,10^. The discovery of these simple relationships in the growth-rate dependence of gene expression control will stimulate further research into the regulatory mechanisms in primary mammalian cells.

This further research will be facilitated by the multiplexing framework introduced here; it enables scaling the analysis of protein synthesis and clearance rates by mass tags, which can further increase the plex and thus the throughput^53,54^. The number of multiplexed samples can also be increased with existing mass tags by modeling the overlap of isotopic envelopes, as done by JMod^55^. Further technological improvements include multimodal mRNA and protein measurements from the same cell and improved estimates of measurement noise, which are essential for estimating the contributions of different regulatory mechanisms^8,56^.

The accuracy and scale of our dataset enabled functional analysis based on covariation of proteins and RNA within a cell type. Advances in single-cell proteomics^57–61^ enable analyzing thousands of single cells^31,62–64^ and performing protein covariation analysis^28,65,66^. Here, we analyze differences between mRNA and protein correlations and their origins. We find that regulated protein clearance strongly contributes to the differences in mRNA and protein covariation patterns (Fig. 6), but it does not account for all differences. In addition to measurement noise, additional contributors include translational regulation and the longer half lives of proteins; thus, transcriptional bursts are integrated into steadier protein levels that reflect the transcriptional history of the cell, not just a momentary point estimate of mRNA abundance^67^.

Differential analysis between young and old mice highlighted significant changes in protein abundance mediated to a substantial degree by changes in protein clearance rates. These effects are strongly cell-type specific, which can help explain the small magnitude of tissue-level protein changes^7^. While the scope of exploration across age groups was limited by the previously mentioned exclusion of the two 24 month mice part of the 10 day metabolic pulse group, the consistent signal across the two replicates suggests cell type specific effects of age on the proteome. These differences show considerable control by changes in protein clearance, in line with expected decline in proteostasis with age^68^.

Our approach opens the door to characterizing gene expression regulation in mammalian tissues and identified potential organizing principles in the role of growth rate. It also demonstrates the feasibility of functionally interpreting protein covariation within individual cell types, that will be amplified by technological developments^69^.

## Resource availability

### Lead contact

Further information and requests for resources and reagents should be directed to and will be fulfilled by the lead contact, Andrew Leduc.

### Materials availability

This study did not generate new unique reagents.

### Data availability

All raw data and search engine outputs can be found on MassIVE with ID: [MSV000098940](https://massive.ucsd.edu/ProteoSAFe/dataset.jsp?task=4b22e2f4fcde424daf16857507045a4c). Single cell mRNA sequencing data was collected as part of Lin et al.^18^ and was previously deposited to the GEO:GSE244215. Additional processed data needed for reproducing the analysis can be found on Zenodo under DOI 10.5281/zenodo.14902833.

[Supplemental data table 1](https://docs.google.com/spreadsheets/d/1drH8TECB4xBLmzcAQj4X1WllUEkUvOzW/edit?usp=sharing&ouid=109814487119977139380&rtpof=true&sd=true)

[Supplemental data table 2](https://drive.google.com/file/d/1jh5UQa6b71mJ9LkuKBTsMNZtEKoonBMH/view?usp=sharing)

[Supplemental data table 3](https://drive.google.com/file/d/1PCkWvgKFTev7dv02D16p_D3YgjHvdtQf/view?usp=sharing)

[Supplemental data table 4](https://drive.google.com/file/d/1Q-v8gST6spW1dw1HTUWARLddWjy5JIOO/view?usp=sharing)

### Code availability

The code used for data analysis and figures is at: github.com/Slavovlab/sc_metabolic_pulse

## Acknowledgments

We thank Georg Wallmann for assistance generating the custom spectral library used for assessing amino acid recycling, Zhixun Dou and Jayaraj Rajagopal for early discussions, and Vladamir Ondruska and Stefan Binder for help operating and maintaining mass spectrometry instruments. The work was funded by an Allen Distinguished Investigator award through The Paul G. Allen Frontiers Group to N.S., a Bits to Bytes award from MLSC to N.S., an NIGMS award R01GM144967 to N.S., a MIRA award from the NIGMS of the NIH (R35GM148218) to N.S, a UH3CA268117 award from NIH to N.S., and an NIA award R01AG092460 to N.S.

## Correspondence

Correspondence and requests should be addressed to

## Author Contributions

**Experimental design**: A.L. and N.S.

**Sample preparation**: A.L., G.S., Y.X.

**Raising funding & supervision**: N.S., A.F.

**Data analysis**: A.L. and N.S.

**Writing & editing**: A.L. and N.S.

## Methods

### Mouse model, handling and metabolic pulse

All mice experiments were performed in compliance with the Institutional Animal Care and Use Committee at Massachusetts General Hospital. Both male and female C57BL/6 mice, either 24-month-old or 4-month-old, were ordered from the NIA. Murine proteomes were metabolically labeled with 13C6 Lysine. Mouse Express L-LYSINE [13C6, 99%] MOUSE FEED kit was purchased from Cambridge Isotopes Laboratories. Mice were first acclimated to the Lys+0 feed for a period of 10 days before switching to the Lys+6 diet for either 5 or 10 days. Mice were fed 5g per day for either 5 days or 10 days before euthanasia. Mice were euthanized with CO2 followed by cervical dislocation. Tissues were harvested post-euthanasia and perfusion with PBS.

### Tracheal single cell suspension generation

Immediately after euthanasia, murine tracheas were dissected and dissociated to single cells in a two-step protocol. Tracheas were first incubated in 20 u/ml papain and 7.5 u/ml DNase I for 30 min at 37 degrees C with rotation, then triturated to release cells. Cells, debris and undissociated tracheal husks were pelleted and resuspended for a second enzymatic digestion in a mix of 350 u/ml collagenase I, 250 u/ml hyaluronidase, 60 u/ml dispase, 3.2 u/ml papain and 75 u/ml DNase I for 20min at 37°C with rotation. After the second digestion, the husks including the cartilaginous scaffold had been dissociated fully. Single cells were filtered through a 100*µ*m filter and red blood cells were lysed using standard RBC lysis buffer. Single cells were then prepared fresh for either single cell proteomics or mRNA sequencing.

Descriptions of the age, gender and feeding time for each mice can be found in the [metadata file](https://zenodo.org/records/14827610/files/Miceotopes_file_annotations.csv?download=1) on Zenodo.

### Single-cell and bulk sample preparation

Mouse trachea single-cell suspensions were collected on 4 different days, resulting in 4 different single-cell sample preparations following the procedure outlined below.

Samples were prepared using the nPOP sample preparation method for multiplexed single cell proteomics^34^ using cell permeability staining to select only intact cells for MS analysis^64^. Cells were concentrated at 1000 cells per *µ*L in 1x PBS and were incubated on ice and in the dark for 20 min with Sytox Green Dead Cell Stain at a concentration of 1 *µ*M (Thermo Fisher S34860). Cells were then washed to remove dye and resuspended in 1X PBS at a concentration of 300 cells per *µ*L for eventual cell sorting. Cells were then sorted in a volume of 300 pL into 9 nL of 100% DMSO droplets on the surface of a fluorocarbon-coated glass slide for cell lysis using the CellenONE cell sorter and liquid handler. Cells sorting gates excluded all cells with measurable green fluorescent intensity (Sytox Green positive) and selected cells between 11 and 27 *µ*m in diameter with elongation below 2. The single cells were incubated for overnight in digestion buffer of 100 ng/*µ*L Trypsin Gold (Promega V5280) and 20mM HEPES. Droplet stability was maintained with the aid of the CellenONE’s humidifier and slide cooling system, set to 75% relative humidity and one degree below the dew point.

The remaining single-cell suspension was used to prepare bulk digest for spectral library generation. Briefly, the remaining cells were pelleted and resuspended in mass spectrometry grade water at a concentration of 1000 cells per *µ*L. The cells were then lysed by freezing to -80 degrees C and then heating to 90 degrees C for 10 minutes^70^. The lysate was digested overnight at 37 degrees C by adding trypsin and 8.5 pH TEAB buffer to a concentration of 20 ng/*µ*L and 100mM, respectively.

The next day, the nPOP single-cell samples were labeled with 20 nL of mTRAQ reagents dissolved in 100% DMSO at a concentration of 1/60th unit per *µ*L. Samples were pooled using the CellenONE in a 50%/50% solution of acetonitrile and water and dispensed into a 384 well PCR plate, dried down in a speed vac and stored at -20C for later injection for LC/MS analysis. Bulk samples were labeled by adding 20 *µ*L of mTRAQ d0 label at a concentration of 1/20th unit per *µ*L to 40 *µ*L of digested bulk peptide at a concentration of 1000 cells of digested peptides per *µ*L. Samples were incubated for 1 hour and then dried down and resuspended in 40 *µ*L of 0.1 % Formic acid and 0.02 % DDM for LC-MS/MS injection.

### LC-MS/MS data acquisition

All samples were separated via online nLC on a Vanquish Neo UHPLC using a 25cm x 75 *µl* IonOpticks Aurora Series UHPLC column (AUR2-25075C18A). Single-cell sets resting in the wells of a 384-multiwell plate were re-suspended in 1 *µ*l of 0.1 % Formic acid, 0.02 % DDM for injection. All samples were run on a 30 minute total length run to run method with 15 minutes of active gradient. 0.1 % formic acid in water was used for Buffer A and a 20% solution of 0.1% formic acid mixed in 80 % acetonitrile was used for buffer B. A constant flow rate of 200nl/min was used throughout sample loading and separation. Samples were loaded onto the column for 20 minutes. The gradient started at 6 % buffer B and ramped to 38% B buffer over 20 minutes. The gradient was then ramped to 95% B buffer over 2 minutes and stayed at that level for 3 minutes to remove excess peptide and protein residue from column. The gradient then dropped to 1% B buffer over 0.1 minutes and stayed at that level for 4.9 minutes for the re-equilibration.

All samples were analyzed on a timsTOF SCP mass spectrometer. Instrument control and data acquisition were performed via timsControl 3.1. The data acquisition method consisted of 8 PASEF frames with 26 Th MS2 windows (1 Th overlaps). An MS1 scan was taken every 2 PASEF frames to increase the frequency of precursor sampling^71^, resulting in 4 MS1 scans per duty cycle. The MS1 scan range was 100-1700 m/z, while MS2 scan range was 300-1000 m/z. The 1/K0 range was between 0.64 and 1.20 and collision energy was set at 20eV at 1/K0 = 0.60 and 59eV at 1/K0 = 1.60, collision RF was set to 2000 Vpp. The ramp and accumulation times were 100 milliseconds and estimated duty cycle time is 1.28 seconds.

### sc-mRNA sample preparation, sequencing, and read alignment

The single-cell suspensions were used for barcoding and library preparation with a Chromium Next GEM Single Cell 3’ Kit v3.1 according to the manufacturer’s protocol. Libraries were sequenced on NovaSeq 6000 S2 with paired-end reads at a depth of 50,000 reads per cell. Reads were processed and aligned to the mm10 mouse genome reference using CellRanger. We collected these data and reported them as part of Lin et al.^18^. The data was previously deposited to the GEO: GSE244215.

### Interpreting raw mass spectra

All MS runs were searched with DIA-NN v1.9.0^6^. The two bulk runs were first searched using an *in silico* generated spectral library from a murine protein sequence fasta downloaded from Uniprot in September 2024 with mTRAQ d0 specified as a fixed modification with mass of 140.0949630177 on all n-termini and lysine. Additional parameters specified were scan window of 5, Mass accuracy of 15 ppm, and MS1 accuracy of 5 ppm. All other parameters were set to defaults. The resulting search generated a empirical spectral library with 4,056 proteins from 56,321 precursors that was used to analyze the single-cells samples.

Single-cell raw files were searched with match between runs in batches of roughly 600 raw files grouped by the days in which the samples were prepared. DIA-NN settings were the same as for the bulk samples with the addition of the peak-translation and channels arguments for plexDIA. Four channels were specified for mTRAQ d0 and light lysine, mTRAQ d0 and heavy lysine, mTRAQ d8 and light lysine, mTRAQ d8 and heavy lysine as follows:

1. mTRAQ,0,nK,0:0
2. mTRAQ,1,nK,0:6.020129
3. mTRAQ,2,nK,8.0141988132:8.0141988132
4. mTRAQ,3,nK,8.0141988132:14.0343278132

We also performed an additional search with the intention of identifying missed cleaved peptides that were generated with both heavy and light lysines to assess amino acid recycling. To do so, we customized to our empirical library to duplicate all entries so we could manually specify light and heavy lysines. For peptides with two lysines, 4 different precursors were created, one with both light, one with both heavy, and the two combinations of one light and one heavy. Runs were then searched with mTRAQ d0 as a fixed modification and two channels for peptides labeled with d0 and labeled with d8. The downside to this approach is that the peak translation would not apply across heavy and light precursors with the same label. Thus the results were only used for comparing the heavy and light distribution for peptides with multiple lysines, subsequently used for determining the rate of amino acid recycling.

### Plotting raw mass spectra

To plot the raw mass spectra, we utilized the alphaTims GUI^72^ to locate precursor ions at the retention time, M/Z, and ion mobility range for peptide LYSEFLGK as reported by DIA-NN.

### SCP pre-processing and batch corrections of abundances

Each single cell protein data set defined by individual nPOP sample preparations was pre-processed using the QuantQC R package^35^. The package first maps relevant meta data to the DIA-NN report file by integrating the dispensing log files from the cellenONE. This includes information such as cell size, label, and run order.

For processing of protein abundances, first, light and heavy intensities for each lysine containing peptides from a given single cell were summed to reflect the total abundance of the peptide. To remove failed cells, total MS intensity summed from all peptides was plotted for all single cells including negative control droplets that received all the reagents but no single cell. An intensity threshold was then set that removed all negative control droplets along with all single cells that had intensities smaller than the negative control with the highest signal. The peptide by single cell matrix was then generated, allowing maximum 5 peptides mapping to the same protein. For proteins with greater than 5 peptides, the most abundant 5 peptides, computed from average intensity across all cells, were chosen. Data was normalized to relative *log*_2_ fold changes via the method described in Khan et al.^31^. Briefly, sample loading normalization was performed using the intersected median across all cells as opposed to that of each cell. Specifically, a reference vector was created from the median value for each peptide across all cells. The median difference between the non missing peptide intensities from each cell and the reference vector were then used to scale all peptide intensities for that cell. Each peptide was then divided by the mean expression across all single cells and the data was log transformed.

Next, to control for variation that occurs across LC runs, a spline was fit to the vector of peptide fold changes and the order to which the MS samples were run. The degrees of freedom for the spline fit were set to the number of single cells with present values divided by 10 with a maximum value of 20. For all spline fits with *R*^2^ greater than 0.1, the variation due to the spline fit was regressed out by subtracting the fit line.

Peptide data was then collapsed to the protein level by taking the median value for all peptides mapping to the same protein within each cell type, resulting in a protein by cell type matrix. To correct for bias due to mTRAQ label used, data was first imputed using KNN with K equal to 3, and then batch corrected using the ComBat R package with the label (d0 or d8) specified as a co-variate.

To generate the final data matrix for all data sets, the data for each sample prep was concatenated together and combat was applied with the sample preparation as a co-variate.

### Clustering, dimensionality reduction and cell type assignment

Cell types for mRNA data were annotated previously as described in Lin et al.^18^ Louvain clustering and UMAP dimensionality reduction was performed on the final protein by cell data matrix via the Seurat R package. Clustering resolution was set to a value of 0.1. Cell types for each cluster were annotated using established cell type markers as shown in Supplemental Fig. 1a. To validate clustering, the average protein and mRNA *log*_2_ fold changes were correlated for all cell types and showed strong agreement for corresponding cell types, Fig. 1c.

### Accounting for missing values in SCP data

To effectively estimate protein abundances at the cell type level, we sought to account for missing data in individual single cells. To do this we first needed to distinguish between missing values due to 1) technical factors such as run to run variability, labeling bias, and variable quantification depth due to cell size variation and 2) reduced abundance due to type specific regulation. This functionality, as described below, is incorporated in the QuantQC package.

To this end, we first formulated a technical effect abundance model that predicted the abundance of each peptide data point based on LC-MS/MS run order, sample preparation batch, and mass tag label. First, we started with the peptide level raw abundance data across single cells, and normalized the data matrix to *log*_2_ relative fold changes based on the reference vector approach described above. We then fit a spline model to each peptide fold change using the run order as the independent variable. The spline was fit separately for samples with either mTRAQ d0 or d8 labels and for each sample preparation batch. Again, the degrees of freedom for the spline was set to floor of the number of data points divided by 10, with a max value of 20. We then used this model to predict abundances for all present and missing peptide data points.

Next, we formulated two different missingness models. Each model was trained to predict whether a peptide data point is present or missing by fitting a logistic regression based on a number of features. The first model predicted missingness due to technical factors by utilizing 1) the abundance prediction from the technical effect abundance model and 2) the cell size. The second model included 3) the assigned cell type as a feature to capture missingness due to cell type specific regulation.

The difference in number predicted missing values between the technical only and cell type logistic regression models was taken as the number of single cells missing due to biological reasons, i.e., the peptide abundance is below the detection limit of the measurements. To better estimate the average protein abundance in a cell type in the presence of biological missingness, the values below the limited of detection were assumed to equal the minimum value measured in this cell type. The cell type averages were then computed including the included minimum values. This correction resulted in improved agreement of fold changes estimated across cell types when compared to scRNA-seq, Supplemental Fig. 2f-h.

### Estimating clearance rates in vivo

To estimate the change in the fraction of light lysine in the unbound lysine pool over time, *γ*(*t*), we first calculated the fraction of heavy lysine in the free pool at our two time points. To do this, we utilized the average ratio of partially labeled to fully labeled peptides as done by Jovanovic et al.^10^. This ratio was computed as the average across all miscleaved peptide from all single cells within each cell type. We then fit an equation of the form *γ*(*t*) = 1 − 0.5*e^−at^* − 0.5*e^−bt^* to estimate the change in available light lysine over time as previously developed and used in ref.^38^ for each cell type individually. Cell types were also fit separately within each age group. Because of the loss of old mice from the 10 day experiment, we estimated the recycling ratio for the old mice at 10 days to be equivalent to the difference between the old and young mice at day 5, Supplemental Fig. 3b.

Having solved for the change in the free lysine pool over time, we then formulated the change in any given light peptide over time 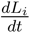 via the equation:

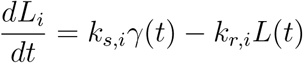

We then integrate this equation to solve for *L_i_*(*t*) for the initial condition *L*(0) = *L*_0_ where *L*_0_ is defined as the sum of heavy and light intensities under the steady state assumption that overall protein concentration is not changing over time. Integrating and further substituting *k_s,i_*via the steady-state equality 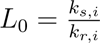. This results in the following equation:

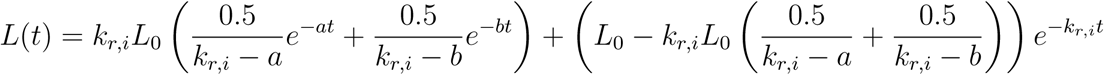

The arithmetic was performed in Mathematica to avoid manual integration errors. Lastly, this equation was then solved for via a non-linear solver to find *k_d,i_* for each peptide from each single cell, with equation parameters for *γ*(*t*) depending on the cell type and mouse age.

### Steady state basal-to-secretory transition analysis

To test the steady-state assumption against a candidate dynamic transition, we constructed a pseudotime trajectory from basal to secretory cells and tested whether newly synthesized heavy proteins predict the future mature (light) proteome. Peptides with fewer than 200 non-missing values across basal and secretory cells were removed, remaining values were normalized by reference-vector log normalization^35^, and residual missing values within each gene were replaced by the gene median for the distance computation. Principal components (PCs) were computed on the resulting gene-gene correlation matrix. A principal curve was then fit to the first 5 PCs with 3 degrees of freedom, and cells were ordered along the resulting curve to define pseudotime, Extended Data Fig. 4a. Marker proteins Krt5 (basal) and Arg2 (secretory) were plotted along pseudotime to validate the trajectory direction, Extended Data Fig. 4b.

To test for a dynamic transition, we compared heavy and light abundance profiles along pseudotime. If cells were actively transitioning between states, newly synthesized heavy protein should lead the mature light protein: for a rising marker such as Arg2, heavy would rise first, and for a falling marker such as Krt5, light would fall first. Cells were binned into 50 equally spaced pseudotime windows, and heavy and light per-protein z-scored abundances were computed for each window, Extended Data Fig. 4d.

To test for localized transitions that a whole-trajectory view might miss, we performed a sliding-window analysis analogous to trajectory-lag comparisons used in single-cell RNA velocity^42^. For each reference position along the trajectory, we defined a reference window of heavy protein abundances centered at that position (window size = 3 adjacent bins). We then correlated this reference heavy profile against the light protein profile at every other position along the trajectory. A dynamic transition would yield higher correlations to future light windows (cells’ current newly-synthesized proteome predicts future mature proteome) than to past light windows. We quantified this asymmetry at each reference position as *θ* = C̅F̅ − C̅P̅, the difference between the average Spearman correlation to the three following windows (CF) and the three preceding windows (CP). Across the trajectory, *θ* averaged 0.01 with no sustained positive regions, Extended Data Fig. 4f, indicating no detectable dynamic basal-to-secretory transition in our data.

### Computing gene regulatory parameters

The following derivations are more comprehensively derived in Baum *et al.*^19^. Briefly, in cells that are not growing, under the steady state assumption protein concentrations can be formulated as a balance of rate of synthesis and degradation.

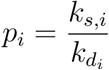

In cells that are growing, the steady state model can be maintained under the assumption that protein concentrations remain constant as cells double the amount of all proteins at an equal rate. However, under this regime, we need to introduce an additional term to reflect protein clearance due not only to degradation, but to dilution as well.

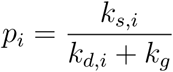

The dilution rate *k_g_*is constant for all proteins and is inversely proportional to the rate of cell division, *T_CC_*, *k_g_* = *log*(2)*/T_CC_*. For both models, we define the overall amount of protein clearance from degradation and dilution is *k_r__i_* = *k_d__i_* + *k_g_*, with *k_g_* = 0 in the absence of growth.

To determine the average cell division time across a population of single cells, we leverage the fact that histones degrade very slowly, but double and partition themselves roughly evenly upon cell division. Thus, the distribution of heavy over light histone abundances takes on a discrete nature where it is possible to asses the number of times a cell had divided. We then use the following equation to determine the doubling time, *N* (*t*) = *N*_0_2*^t/TCC^* where *t* is the pulse duration, *N* (*t*) is the number of single cells in the population at the pulse duration *t*, and *N*_0_ is the initial number of cells. *N*_0_ can be computed as the following:

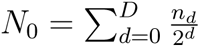

where *n_d_* are the number of cells that have divided d times. Error bars are the standard deviation of *T_CC_*estimates across the 4 sample preparations.

Lastly, the synthesis rate can be further decomposed into the number of mRNA copies for the i*th* gene in the cell, *mRNA_i_*, multiplied by the average rate the mRNA is translated.

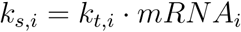

Resulting in the final expression defining the regulation of the *ith* gene:

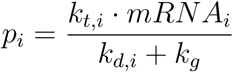

### Handling measurement noise for absolute abundance control

To handle measurement noise when estimating the extent that mRNA abundance, translation, and protein clearance influence absolute protein abundance, we utilized reliability measurements from average estimates from correlating random subsets of single cells. For two random subsets of cells, we required at least 5 observed values to contribute to each average before computing the reliability. From this approach, we are underpowered to estimate the influence of systematic biases contributing to absolute copy number of proteins and mRNAs. Phenomena such as ionization and digestion bias for protein measurements, and cell lysis, transcript capturing and PCR biases for mRNA measurements might bias absolute abundance estimates across different proteins and transcripts. Thus, estimates of reliability for absolute protein and mRNA abundances were multiplied by 0.7 to reflect these systematic biases,^12,73^. This adjustment reflects comparison of different mRNA sequencing technologies and peptides generated from different protease digests. No further correction was applied to the protein clearance estimates as these biases should cancel out when computing the ratios between heavy and light precursors.

We then used the Spearman correction to correct the percent variance explained by both clearance and mRNA abundance^8^. In the corrected estimates, the translation contributes the remaining variance. In the uncorrected estimates, the raw *R*^2^ of each modality is displayed in Supplemental Fig. 7e. Estimates made from the remaining variance correlate well with those made by regressing the translation rates against protein abundance, *ρ* = 0.75, Supplemental Fig. 7d.

To estimate uncertainty in the variance-explained statistics, we applied a nonparametric bootstrap procedure at the level of genes averages. For each cell type, paired values of the regulatory variable (e.g., clearance rate) and protein abundance were resampled with replacement to the same size as the original dataset. The variance-explained statistic was recalculated for each bootstrap sample, yielding a distribution of bootstrap estimates. We repeated this procedure 1,000 times per cell type and summarized the dispersion of the bootstrap distribution by its standard deviation, which we report as the bootstrap error for the variance-explained measure.

### Estimating relative levels by Bayesian inference

To estimate mRNA abundance, translation rate, protein clearance and relative protein abundance of each protein in each cell type, we relied on replicate estimates from peptides mapping to the same protein, individual single cells, and measurements made from sample preparation replicates. We formulated the problem using Bayesian inference to estimate measurement noise for individual quantities in individual cell types.

Specifically, we modeled the log transformed parameters, including relative fold changes compared to the average across cells for protein abundance (*logP*), mRNA abundance (*logM*), translation rate (*logT*), and clearance rate (*logC*). These values were each inferred for for gene *g* in cell type *c* via the log transformation of the previously established relationship between the quantities:

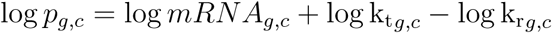

In our model, the protein abundance, mRNA abundance, and clearance rates were the only observed values. Translation rate is inferred from the remaining difference in protein levels not explained by mRNA abundance and protein clearance.

The observed quantities were modeled using Gaussian likelihoods with standard errors derived from the number of replicate observations. For protein abundance (*p̄_m_*) and clearance rates(*c̄_m_*), observed quantities were the peptide level cell type average of the relative *log*_2_ fold changes compared to the average value across cell types for one of the four sample preparation replicates.

Protein abundance observations were modeled as:

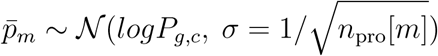

where *g* and *c* denote the gene and cell type indices for observation *m*, and *n*_pro_[*m*] is the number of cells where the peptide was observed in the given replicate. Similarly, observed clearance rates were modeled as:

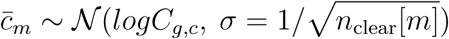

For observed mRNA (*M̄_m_*) quantities were similarly the cell type average of the relative *log*_2_ fold changes compared to the average fold change within replicate.

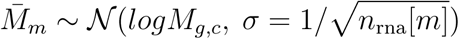

Here *n*_rna_[*m*] was the number of non-zero observations of the transcript in the given cell type and replicate.

We placed zero-centered Gaussian priors with standard deviation that reflected our prior certainty for each quantity:

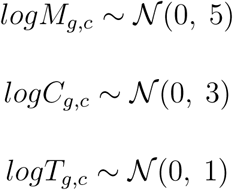

This model enabled us to propagate measurement noise and quantify uncertainty in the estimates of gene expression regulation for each gene-cell type pair, allowing robust decomposition of protein abundance into transcriptional, translational, and degradative components.

### Common protein/mRNA principal component space

To align the protein and mRNA measurements of basal cells along a shared axis of variation, Fig. 3, we constructed a common PC space as performed in Specht et al.^50^ Briefly, we first identified a shared gene set whose covariation structure was concordant between the two modalities. For each gene, we computed its correlation vector across all other genes at both the protein and mRNA level, and correlated the two vectors. Genes whose protein and mRNA correlation vectors agreed at Pearson *r >* 0.4 were retained, resulting in a set of concordantly-covarying genes. The PC basis was then learned from the protein data. Specifically, the gene-gene correlation matrix across the 1,345 basal cells with protein measurements was computed and its eigenvectors used as the rotation matrix. The protein data were centered and scaled per gene and projected onto this rotation to obtain protein PC scores. The mRNA matrix, restricted to the 10,000 basal cells with mRNA measurements and to the same gene set, was then centered and scaled using the protein-derived per-gene means and standard deviations and projected onto the same rotation to obtain mRNA PC scores in the identical basis, Fig. 3a. Any residual missing values in the scaled matrices were set to zero prior to projection. This ensured that the two modalities were expressed in the same coordinate system without either modality biasing the PC directions.

To quantify continuous trends along this axis, cells from both modalities were binned into 10 equal-size bins defined by quantiles of the pooled protein and mRNA PC1 scores. For each bin, per-gene bin averages were computed independently for each modality and were used for the downstream trend correlations shown in Fig. 3b–f. Bootstrap standard errors for each bin statistic were computed by resampling genes with replacement 200 times per bin.

### DepMap CRISPR dependency comparison

To externally anchor the within-basal-cell protein and mRNA trends against proliferation-associated essentiality, we compared per-gene trend correlations to CRISPR gene-effect scores from the DepMap Public release, accessed via the Bioconductor `depmap` package^45,46^. Because DepMap uses human HGNC symbols, mouse gene symbols were mapped to their upper-case human orthologs. For each gene, the median dependency score across all screened cell lines was used, so that more negative values reflect stronger essentiality for proliferation.

For each gene shared between the DepMap panel and our basal-cell dataset, we computed a bin-trend correlation with total protein synthesis. Total protein synthesis for each cell was estimated as the average heavy-lysine incorporation of the top 100 most abundant proteins. Bin-averaged synthesis was computed as the mean of this per-cell quantity within each PC1 bin. For each gene, protein and mRNA bin averages were separately correlated against bin-averaged total synthesis, yielding a protein bin-trend correlation and an mRNA bin-trend correlation per gene. These pergene correlations were then compared to the DepMap median dependency score, Fig. 3f. Genome-wide, protein bin-trend correlations aligned with DepMap dependency at Pearson *r* = −0.25 versus *r* = −0.10 for mRNA bin-trend correlations. For the focused analysis of glycolytic enzymes, the same procedure was applied to a curated set of glycolysis and lactate metabolism genes.

### Correlation analysis across cell types

To determine the extent to which each protein is regulated by mRNA abundance, translation, and clearance, we correlated proteins abundances to the extent of regulation by each mode across cell types. So that a positive correlations would reflect strong regulation for each mode, we utilized clearance half life, *log*(2)*/K_r_*, instead of clearance rate. To determine functional groups of proteins primarily regulated by a single mode of regulation, we performed a gene set enrichment analysis on the correlation distribution for each mode of regulation. Within each mode, we compared the distribution correlation for genes in each gene set with at least 4 proteins present, to the remaining distribution of correlations using a t-test. We then corrected the distribution of p-values within each mode of regulation for multiple hypothesis testing with the Benjamini-Hochberg (BH) method.

### Cell type specific gene regulation analysis

To plot the regulatory influences for gene products that were predominantly regulated by a single mode of gene expression in individual cell type, we plotted in Fig. 4 all genes where the contribution from a single mode was greater than the contribution of the other two. We clustered data individually for mRNA, translation, and clearance and plotted the genes passing this threshold stacked for each modality, Fig. 4f.

To identify functional sets of proteins that were over represented in the genes primarily regulated by a single mode of gene expression, we utilized the compareCluster function from the R package clusterprofiler. Briefly, this utilizes a hyper-geometric test to compare the genes that passed the threshold to the background set of all genes where mRNA, translation, and clearance were quantified. All plotted terms met a significance threshold of FDR of 5 % across at least one of the 6 cell types.

### Single cell protein-mRNA-clearance covariation analysis

We plotted all pairwise correlations for intersected sets of proteins and mRNAs where the proteins had over 100 pairwise observations to robustly support the computed correlation. Curate a list of protein pairs that that directly interact we took all pairwise protein combinations from complexes in the Corum protein data base (release 5.0) as well as including all reported interactions in the STRINGdb with confidence score above 0.7.

To identify protein or mRNA specific correlation modules, we filtered for proteins that contained at least one pairwise correlation above a certain threshold. We first set the threshold at 0.3 for proteins and then plotted the heatmap via hierarchical clusters. We then extracted the clusters at a resolution of the top 7 clusters. Lastly we looked for GO term over enrichment for proteins in these clusters compared to the background set using the clusterProfiler r package (Release 3.21)^74^.

Several terms were enriched for each of the 7 clusters and the most significant GO term was displayed below the heatmap. The analogous procedure was applied to the mRNA data but few transcripts displayed pairwise correlations above 0.3. Reducing the threshold to 0.20, we identified 5 clusters with greater than 10 transcripts. However, only 3 of the 5 clusters contained GO terms with significant over enrichment.

To quantify the contribution of protein clearance to the differences between the corresponding protein-protein and mRNA-mRNA correlations, we started by estimating protein regulation by clearance, *r_c_*; *r_c_* was estimated by correlating protein abundances across single cells to the paired clearance rates across the same single cells. We then plotted the difference between pairwise correlations for the same pair of gene products at the mRNA and protein level versus the average *r_c_* for the protein pair. The resulting correlation, *ρ* = −0.44, suggests significant contribution of clearance to the difference in correlations.

We then created a linear model including as additional explanatory variables the average absolute half life of the two proteins as well as the average raw counts of each transcript. This increased the explanatory power of the model, R = 0.55.

### Single-cell functional covariation analysis

We test for enriched correlations in functional groups of genes within cell type at the protein and mRNA levels. We performed this analysis in the space of intersected proteins and mRNA quantified in each cell type. For each GO term with at least 4 genes, we performed a t-test between the distribution of all pairwise genes from the go term and the background distributions of pairwise correlations for all other proteins or transcripts. We then corrected the distribution of p-values for multiple hypothesis testing with the Benjamini-Hochberg (BH) method. We plotted all terms which were significant at 1% FDR in at least one cell type at the protein or mRNA level.

### Fibroblast vs basal covariation analysis

To compare within cell type covariation patterns across fibroblast and basal cells, we first computed pairwise correlations for all proteins across single cells within a cell type. For each cell type, we defined correlation vector for each protein as the vector of correlations to all other proteins. To compare covariation patterns across cell types, we correlated these correlation vectors for shared proteins between basal and fibroblast cells. A null distribution was computed by randomizing abundances across cells and proteins for each cell type and computing the same procedure.

We then took a threshold of all proteins with a correlation vector similarity of Pearson correlation above 0.5 and denoted this the set of proteins with common covariation patterns as set 1. The similar correlation structure of pairwise proteins can be observed via the heatmap, Fig. 6c. Clusters were annotated for functional groups with the procedure described for the protein-mRNA comparison, but was only applied on the basal cells due to the existing high correlation similarity between the two cell types. All proteins with a correlation of correlation vector below 0.1 were denoted as set 2 proteins. Differences in the covariation between set 1 and set 2 proteins across cell types were identified by clustering the correlation matrix defined by correlations between all set 1 and set 2 proteins in basal cells, Fig. 6d.

## Extended Data Figures

**Extended Data Fig. 1.**
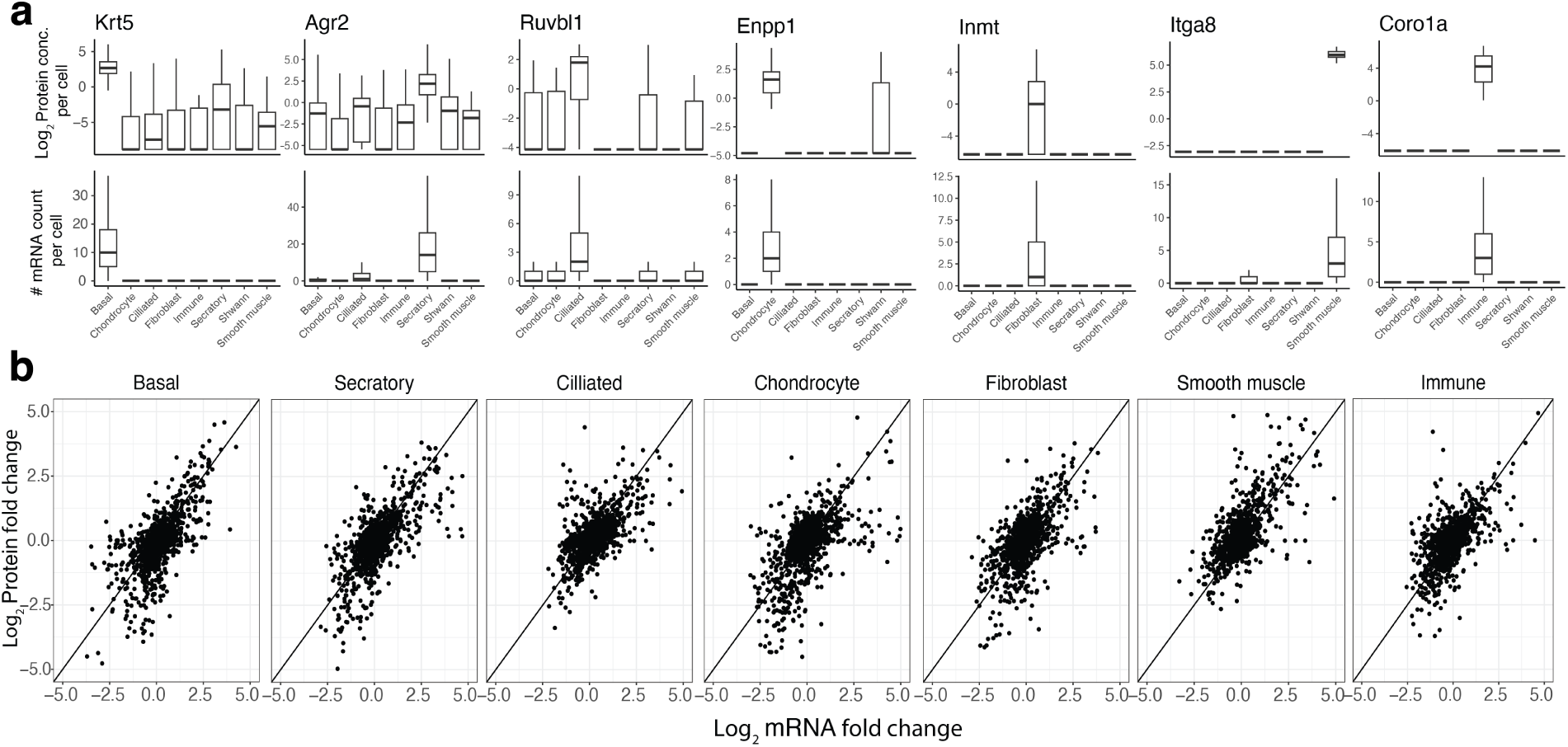
Validating the alignment of the mRNA and protein cell-type clusters. **a**,, Abundance of cell-type specific markers for each of the 7 cell types. The protein levels are plotted as *log*_2_ concentrations, and the mRNA levels as raw counts. **b**, Scatter plots for relative protein levels in each cell type. The levels are plotted as *log*_2_ ratios relative to the mean across all cells types.

**Extended Data Fig. 2.**
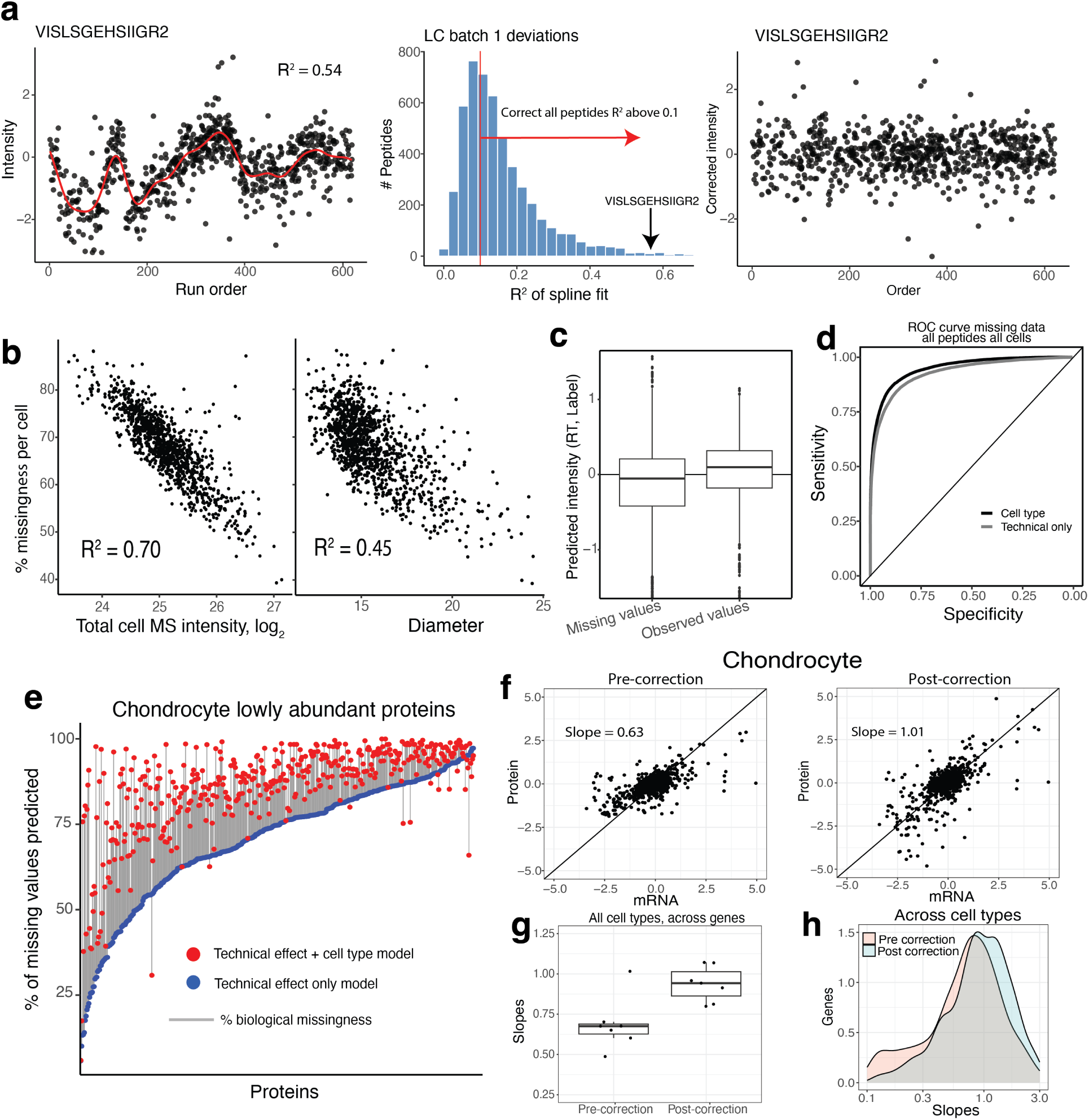
Processing and normalization of the single-cell proteomics data. **a**, Visualization of bias in peptide abundance based on deviations that occur systematically across runs and spline fit to data. *R*^2^ of the spline fit for all measured peptides. Regression of trend was applied to peptides with an *R*^2^ *>* 0.1. The abundance of the peptide VISLSGEHSIIGR2 does not vary with run order after correction. **b**, Fraction of missing peptide data points in a cell is a function of total MS signal measured or similarly the cells and the cell size. **c**, Results of a logistic regression model fit to predict intensity based on retention time and label of the single cell show moderate predictive power on determining whether a peptide data point is missing or observed. **d**, The ROC curve for a logistic regression model fit to predict missingness of peptide data points based on technical factors (cell size, retention time and label). Including the cell-type identity improves the predictions. **e**, For the least abundant proteins in chondrocytes, including cell-type information increased the prediction accuracy for missing values. **f**, *Log*_2_ protein and mRNA fold changes for chondrocytes plotted before and after missing data correction. **g**, Slopes of mRNA protein fold changes across gene products for all cell types plotted before and after correction. **h**, Similar to panel g, but the slopes are estimated across cell types.

**Extended Data Fig. 3.**
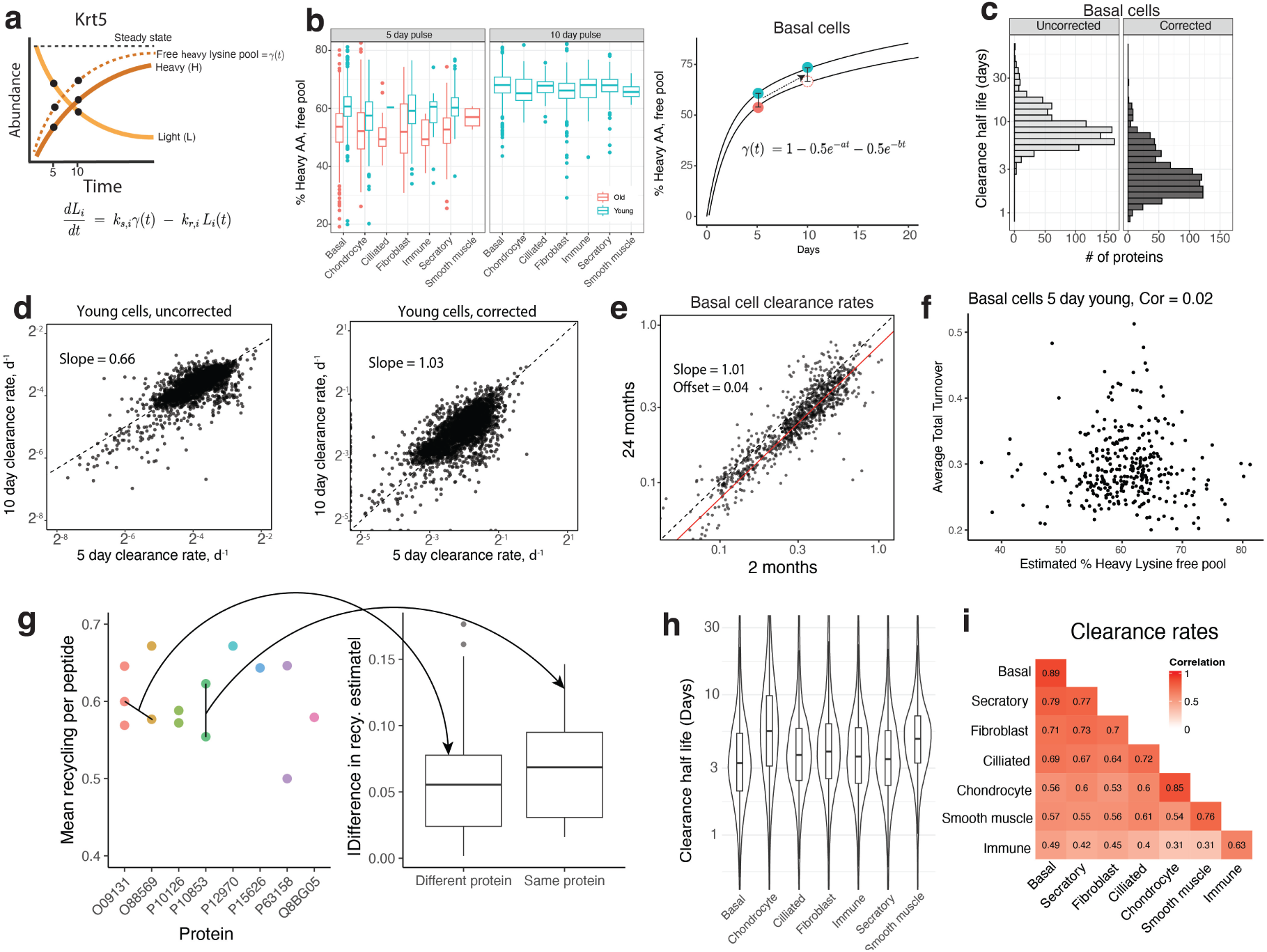
Adjusting clearance for amino acid recycling. **a**, At each time point, three data points are sampled, the heavy amino acid intensity, the light amino acid intensity, and the fraction of heavy lysine available as predicted from miscleaved peptides. **b**, Free heavy lysine pool computed from peptides with missed cleavages as the ratio (HL/HH)/(HL/HH+2). Data points represent average estimates across all peptides for a given single cell. Fit line to the average data points within cell type for 5 and 10 days for the sum of two exponential for basal cells. Due to the loss of old mice at the 10 day time point, the 10 day time point was estimated using the same difference between old and young at the 5 day time point. **c**, Adjusted clearance half-life distribution computed before and after recycling correction. **d**, Slope between 5 and 10 day cells computed before and after the adjustment for amino acid recycling. **e**, Basal cell clearance rate for all proteins plotted for 24 month and 4 month mice. **f**, Estimate fraction of total free lysine that are heavy plotted against the average turnover across all proteins for each cell. **g**, Average heavy lysine fraction across all cells estimated for different peptides grouped by protein identity. Plotted are all pairwise differences in heavy lysine between all peptides plotted by whether the two peptides share or do not share a parent protein. **h**, Clearance rate half-life distributions for all cell types. **i**, Correlation between average clearance rate distributions for all cell types. The diagonal represents the correlation of average clearance rates for the same cell type computed from non-overlapping subsets of single cells.

**Extended Data Fig. 4.**
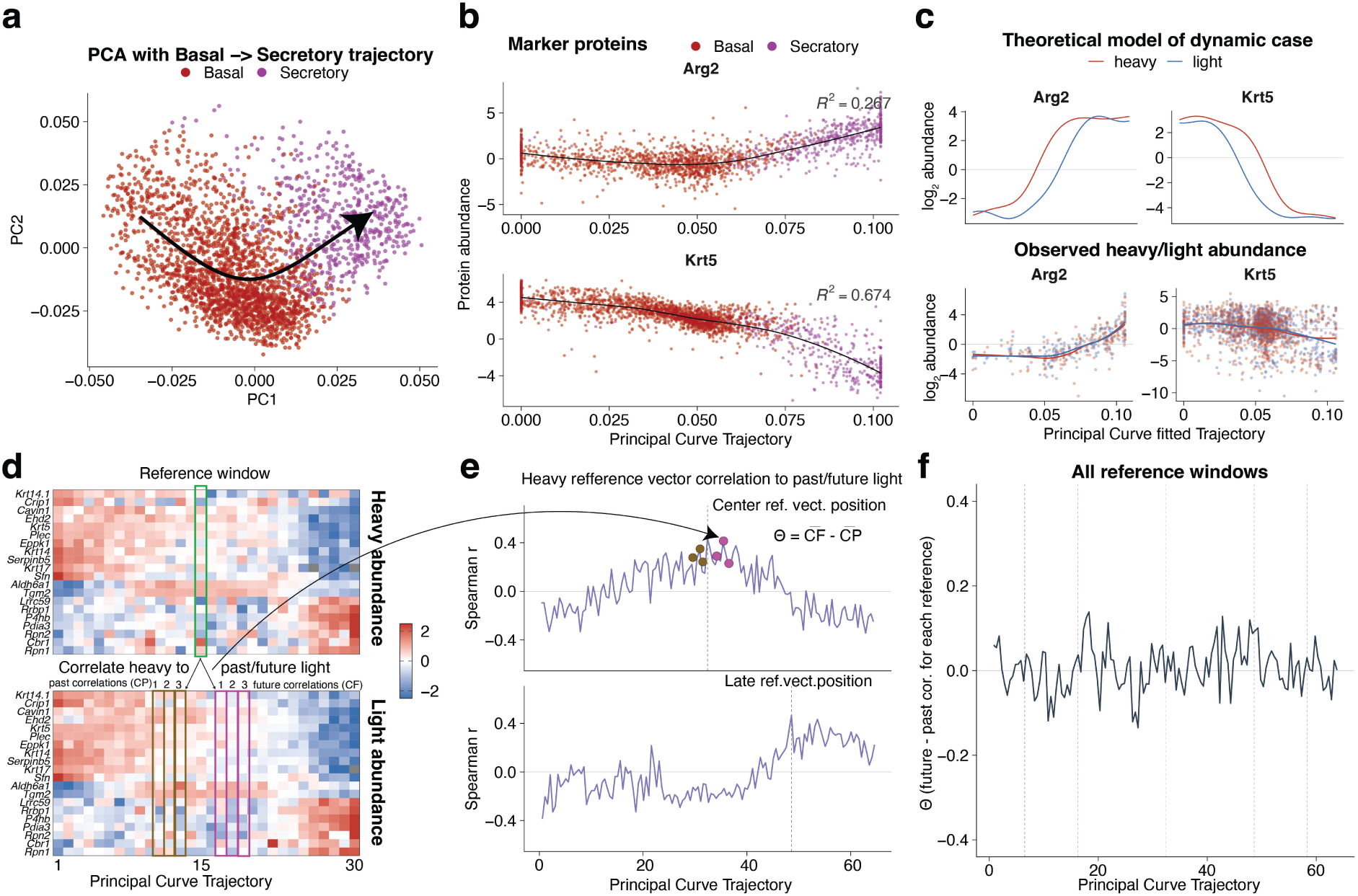
Evaluating separation of time scales and the quasi steady state assumption. **a**, A principal curve fit with 3 degrees of freedom traces a plausible differentiation trajectory across the low-dimensional projection of basal and secretory cells. **b**, The basal marker Krt5 and the secretory marker Arg2 show the expected monotonic decrease and increase along the differentiation trajectory. **c**, The theoretical heavy and light protein traces for a dynamic transition are shown on the top while the empirical ones are shown on the bottom. **d**, Heatmaps of the 20 most variable proteins along the trajectory, binned into 50 equally-spaced windows. To test whether newly synthesized heavy proteins predict the future light proteome, we correlated each reference heavy window to the three preceding and three following light windows. **e**, For an example reference position along the trajectory, correlations between the heavy reference window and light windows at all other positions along the trajectory are shown. We define *θ* as the difference between the average correlation to the three following windows (CF) and the three preceding windows (CP). **f**, The parameter *θ* evaluated at every position along the trajectory.

**Extended Data Fig. 5.**
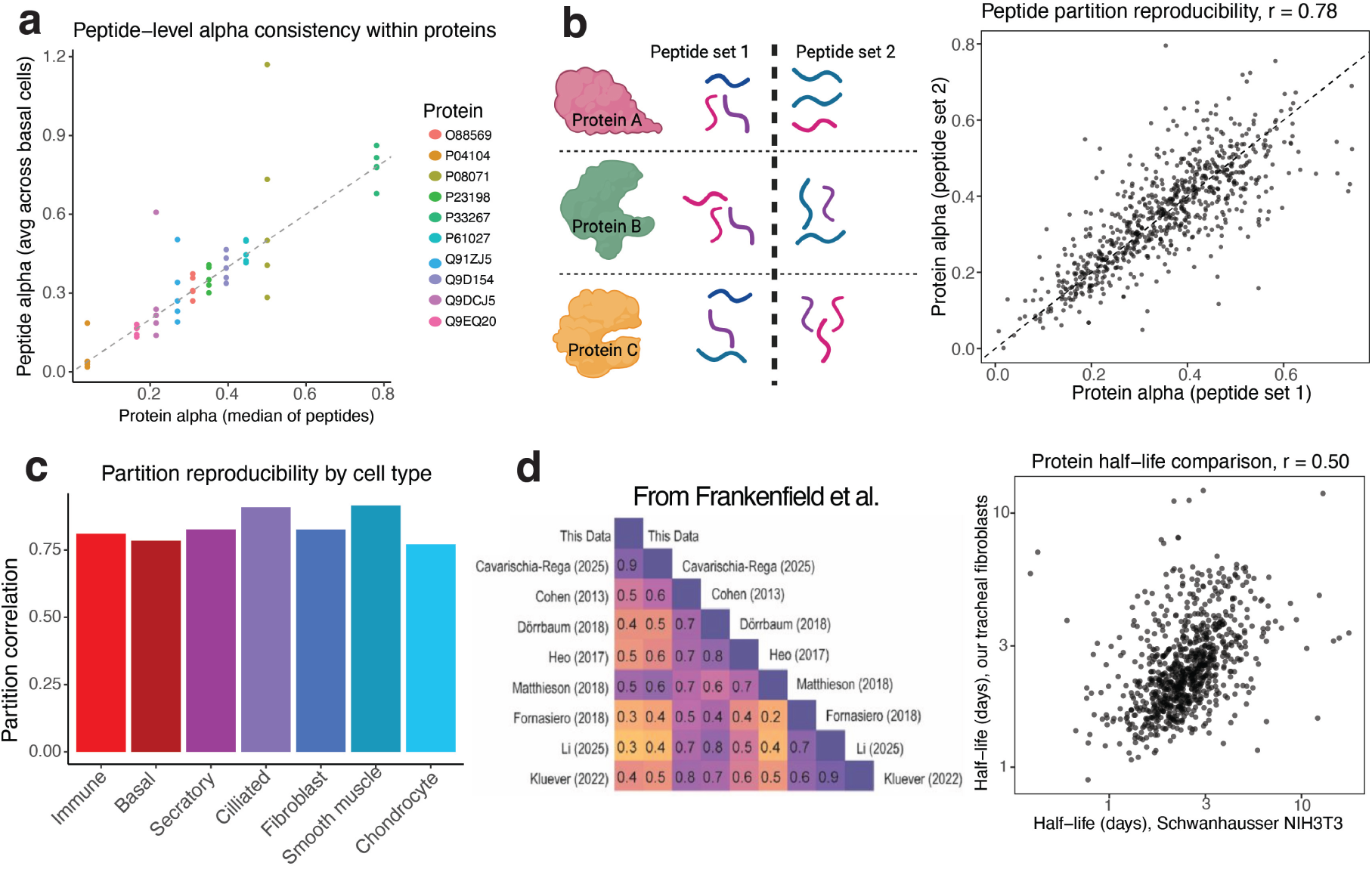
Clearance estimate validation. **a**, For a representative set of 10 proteins, clearance estimates from individual peptides are plotted against the mean estimate for the parent protein. **b**, To assess accuracy of clearance estimates proteome wide, we created two sets of randomly split peptides from all proteins with at least 2 peptides. We then compared clearance values from peptides computed from each set for basal cells. **c**, Clearance estimates from peptide set 1 and set 2 for each cell type. **d**, Frankenfield et al.^44^ compared the similarity of clearance rates measured by metabolic pulse labeling across 9 different studies both in cell lines, primary cells and primary tissues. The clearance rates for primary mouse fibroblasts in our study correlate well (Pearson *r* = 0.50) to previously estimated clearance rates from an in vitro cultured fibroblasts^9^.

**Extended Data Fig. 6.**
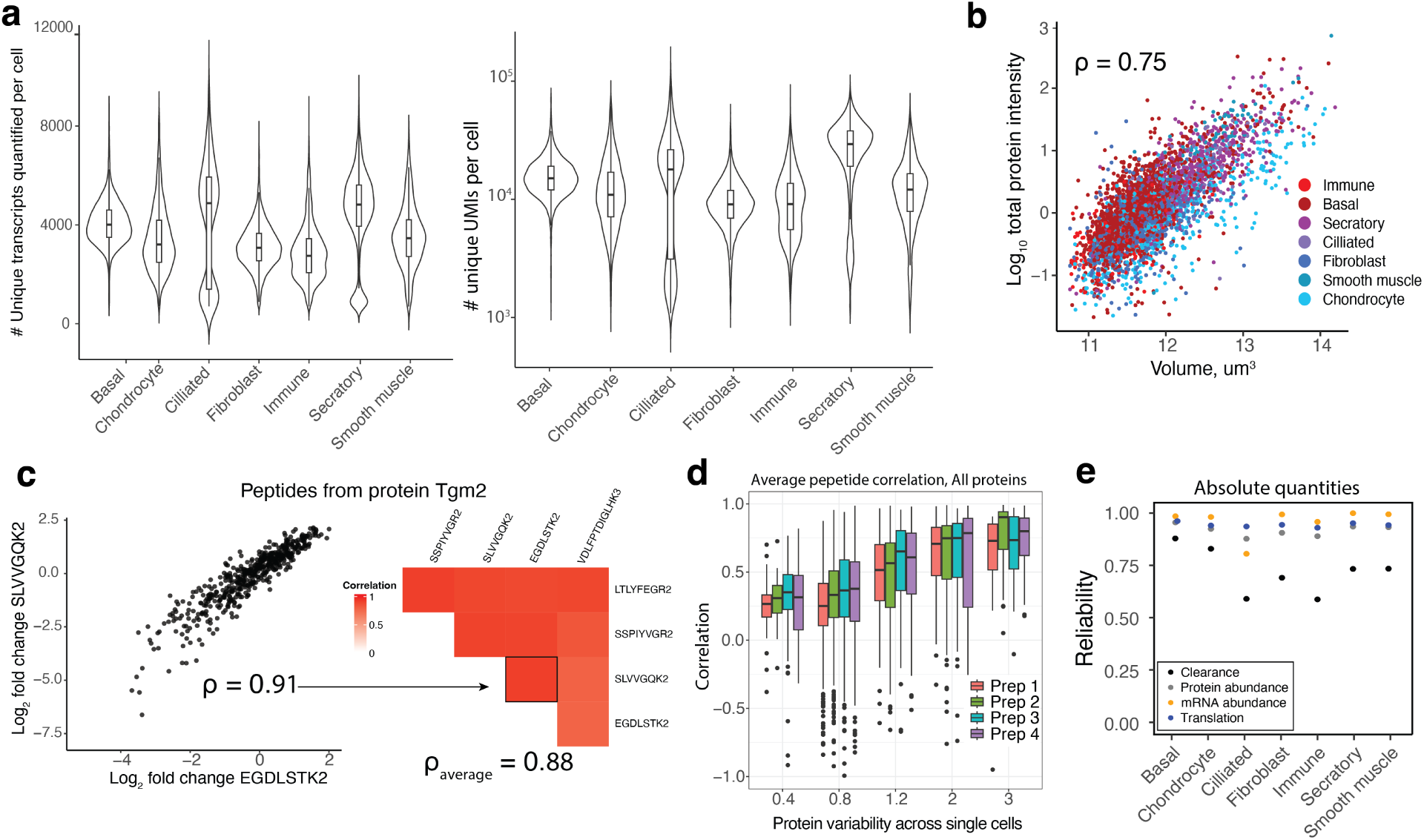
Quantitative accuracy and reliability of mRNA abundance, protein abundance, and clearance rate estimates. **a**, Quality plots for single-cell mRNA data show the number of unique transcripts detected on average per cell and the number of unique UMIs detected. **b**, The log transformed sum of all protein intensity per cell plotted against the cell volume show that total protein is highly proportional to cell size. **c**, Relative protein *log*_2_ fold changes for two peptides from protein Tgm2 across single cells. Correlations of protein fold changes for the to 5 most abundant peptides have an average Pearson correlation of 0.88. **d**, The average correlation for all proteins is plotted against the average protein fold change across single cells for each data set. More variable proteins with higher average fold change correlate more highly. **e**, The average reliability for each modality calculated by taking averages from two random subsets of single cells and plotting the correlation between subsets.

**Extended Data Fig. 7.**
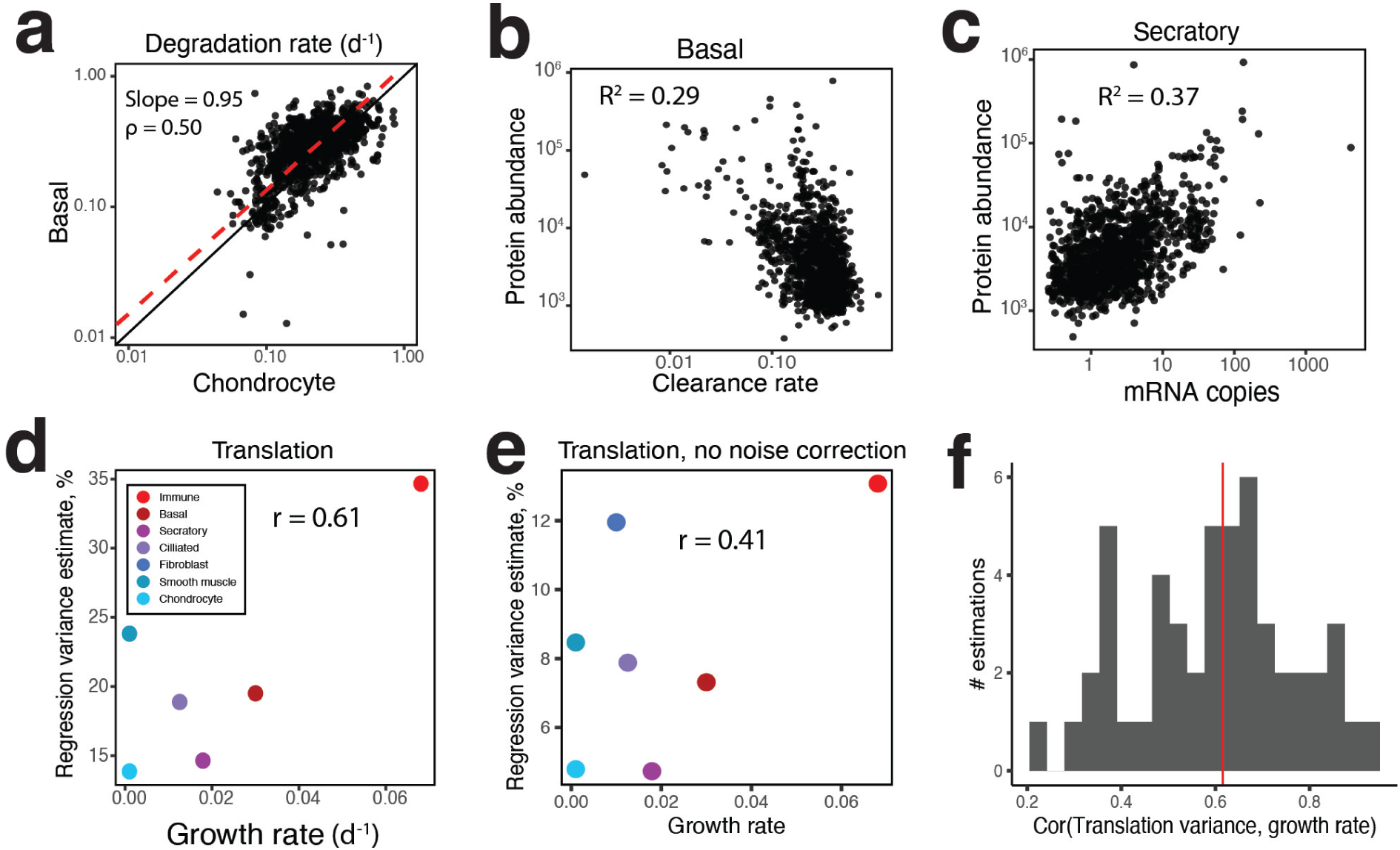
Cell growth modulates regulation of absolute abundance. **a**, Despite the difference in the slope of clearance rates between basal cells and chondrocytes (shown in Fig. 2), the degradation rate slope is 0.95. This suggests that the shallow slope is due to the influence of cell growth. **b**, Scatter plot of clearance rates and absolute protein abundance in basal cells shows *R*^2^ of 0.26. **c**, Scatter plot of mRNA copies and absolute protein abundance secretory cells show *R*^2^ of 0.51. **d**, Scatter plot translation contribution estimates computed from the squared Pearson correlation between translation rates and protein abundance, and cell growth rate still shows strong positive trend. **e**, The same comparison as in panel d, but without any noise correction applied. **f**, Distribution of Pearson correlations between the percent variance explained by translation and cell growth rate obtained from the sensitivity analysis.

**Extended Data Fig. 8.**
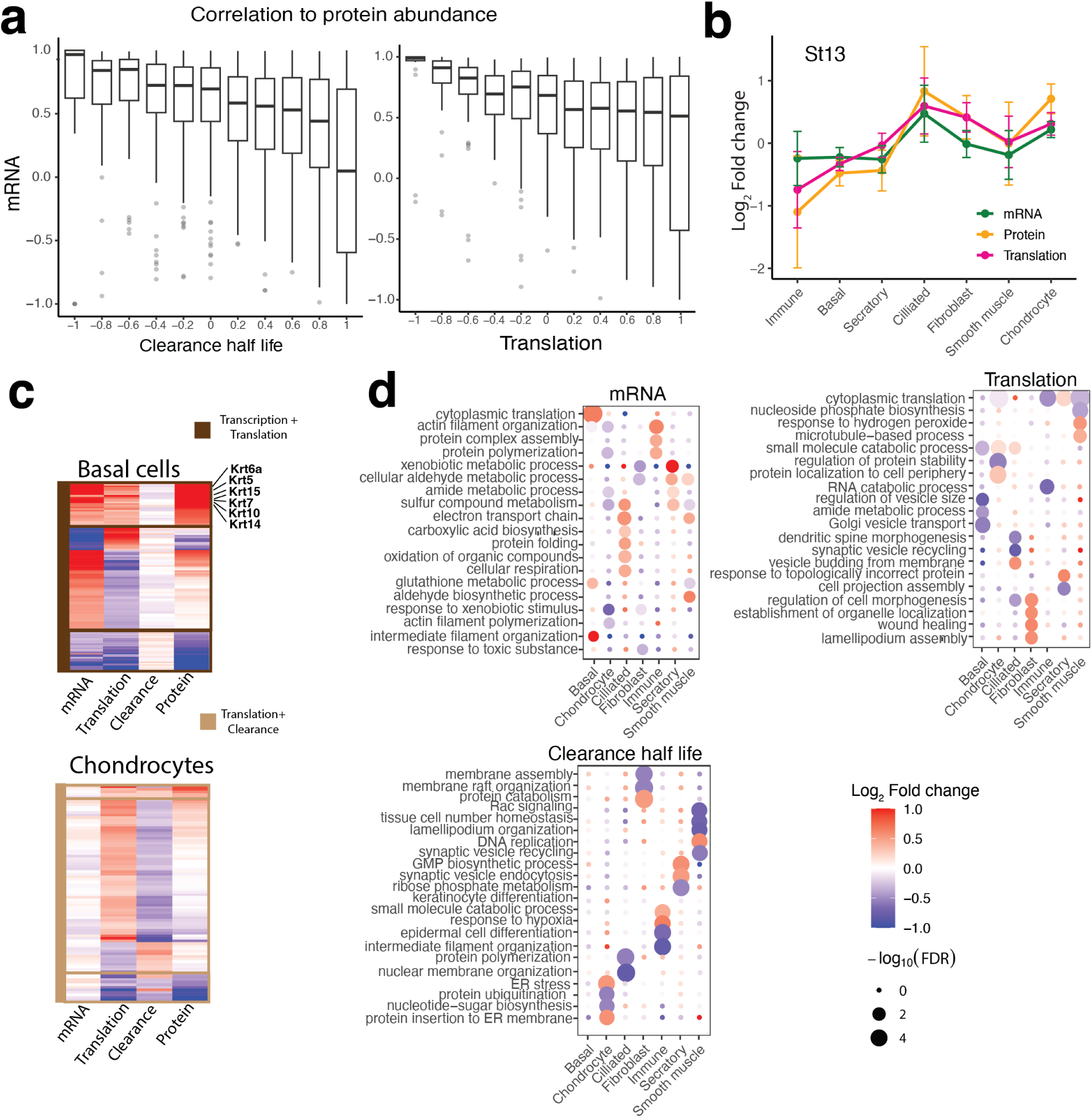
The modes controlling relative protein levels can be shared across cell types or cell-type specific. **a**, Joint distributions of correlations. The top panel shows distributions of correlations between protein and mRNA abundance versus bins of the correlations between protein abundance and protein clearance. The bottom panel shows distributions of correlations between protein and mRNA abundance versus bins of the correlations between protein abundance and translation rate. **b**, Examples of synergistic regulation of St13 across cell types. **c**, A heatmap showing synergistic regulation by mRNA abundance and translation in basal cells, and translation and clearance in chondrocytes. **d**, GO terms for proteins enriched to be regulated primarily through transcription, translation and clearance show cell type specific trends.

**Extended Data Fig. 9.**
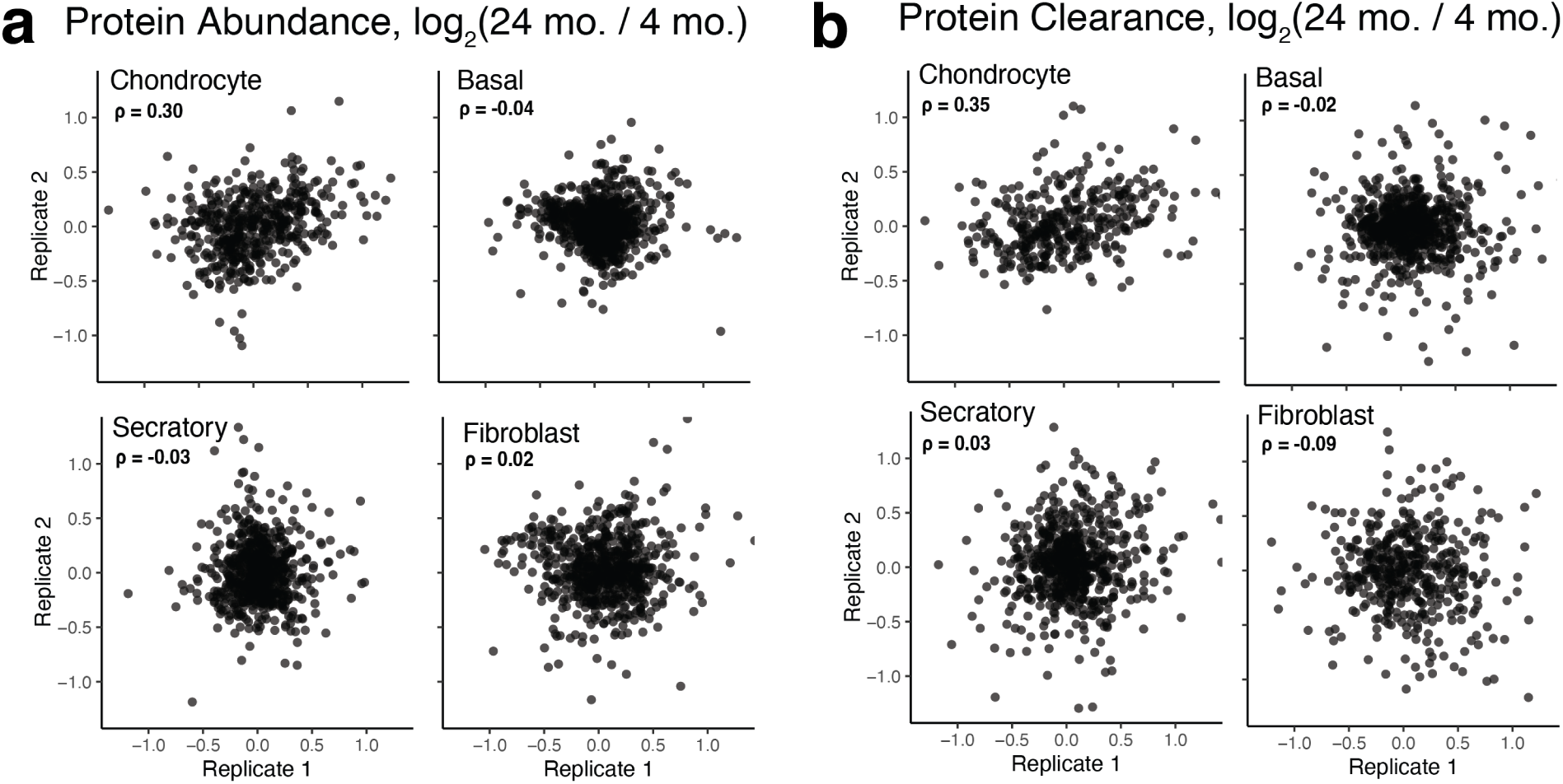
Reproducability of cell type specific aging regulation. **a**, Comparison of protein abundance fold changes between old (24 month) and young (4 month) old mice for replicates experiments. **b**, Comparison of protein clearance fold changes between old (24 month) and young (4 month) old mice for replicates experiments.

**Extended Data Fig. 10.**
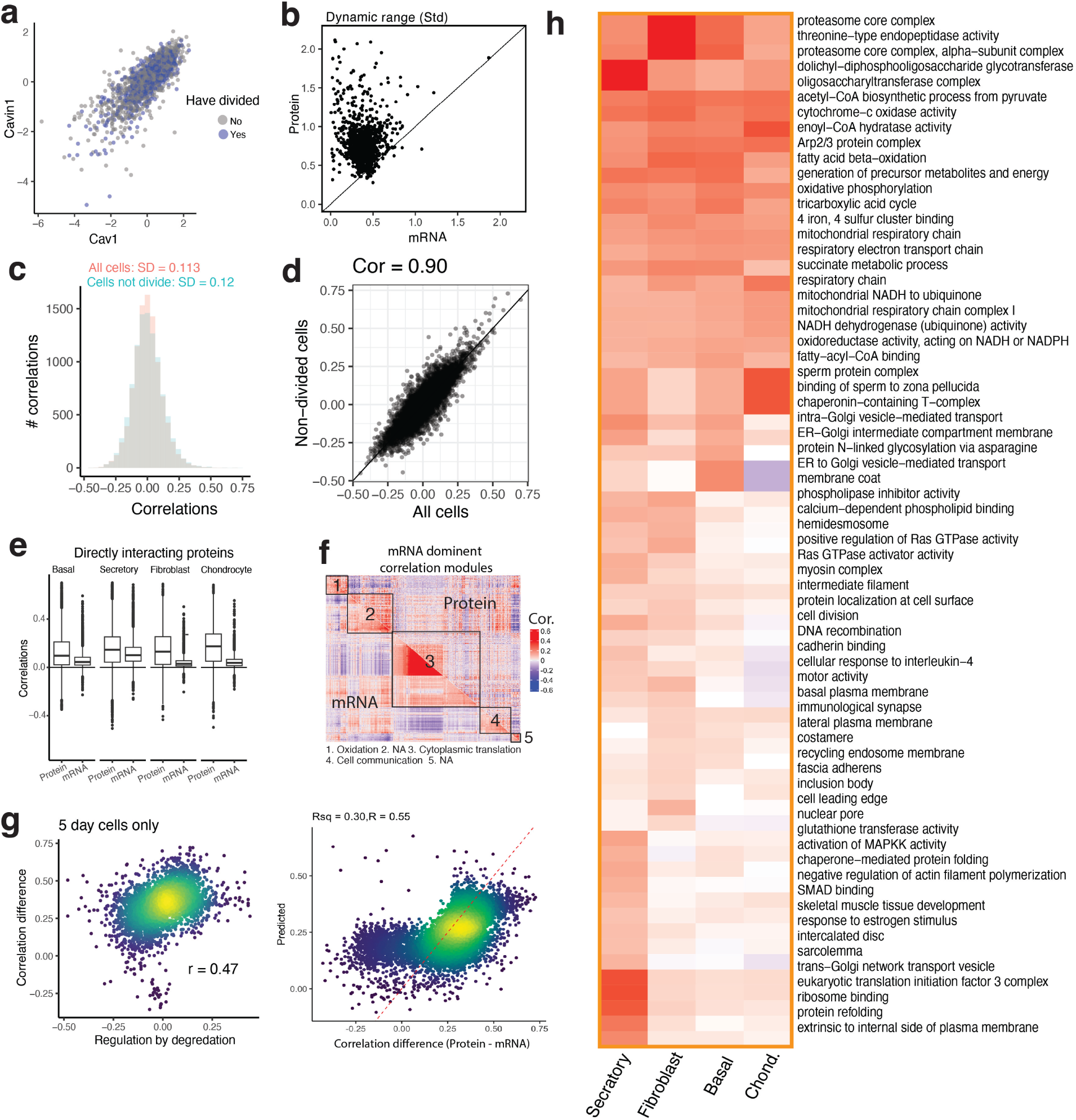
Single-cell mRNA and protein covariation. **a**, Scatter plot showing the correlation between Cavin1 and Cav1. Colors represent whether a cell has divided in the time that has passed since pulse labeling started. **b**, A comparison of the dynamic range of mRNA and protein abundance within basal cells indicates larger variation for proteins. The dynamic range is estimated by the standard deviation of *log*_2_ fold changes relative to the mean for protein and mRNA levels. **c**, Distribution of pairwise protein-protein correlations computed across all basal cells (pink) and across only basal cells that have not divided during the pulse (green). **d**, Scatter of pairwise protein-protein correlations computed in all basal cells against the correlations computed in the subset of non-divided cells. **e**, Correlations between directly interacting proteins within each cell type for proteins that have over 100 pairwise observations and the corresponding mRNAs. **f**, mRNA dominant correlation modules lack enriched GO terms and exhibit low correspondence to protein data. **g**, Prediction of the difference between protein-protein and mRNA-mRNA correlations by a linear model that incorporates the protein clearance rates, the correlation between the relative clearance rates and proteins levels, and the average mRNA counts. **h**, A heat map displaying average correlations for the full list of GO terms that were significant in at least one cell type.

## Supplemental Figures

**Supplemental Fig. S1.**
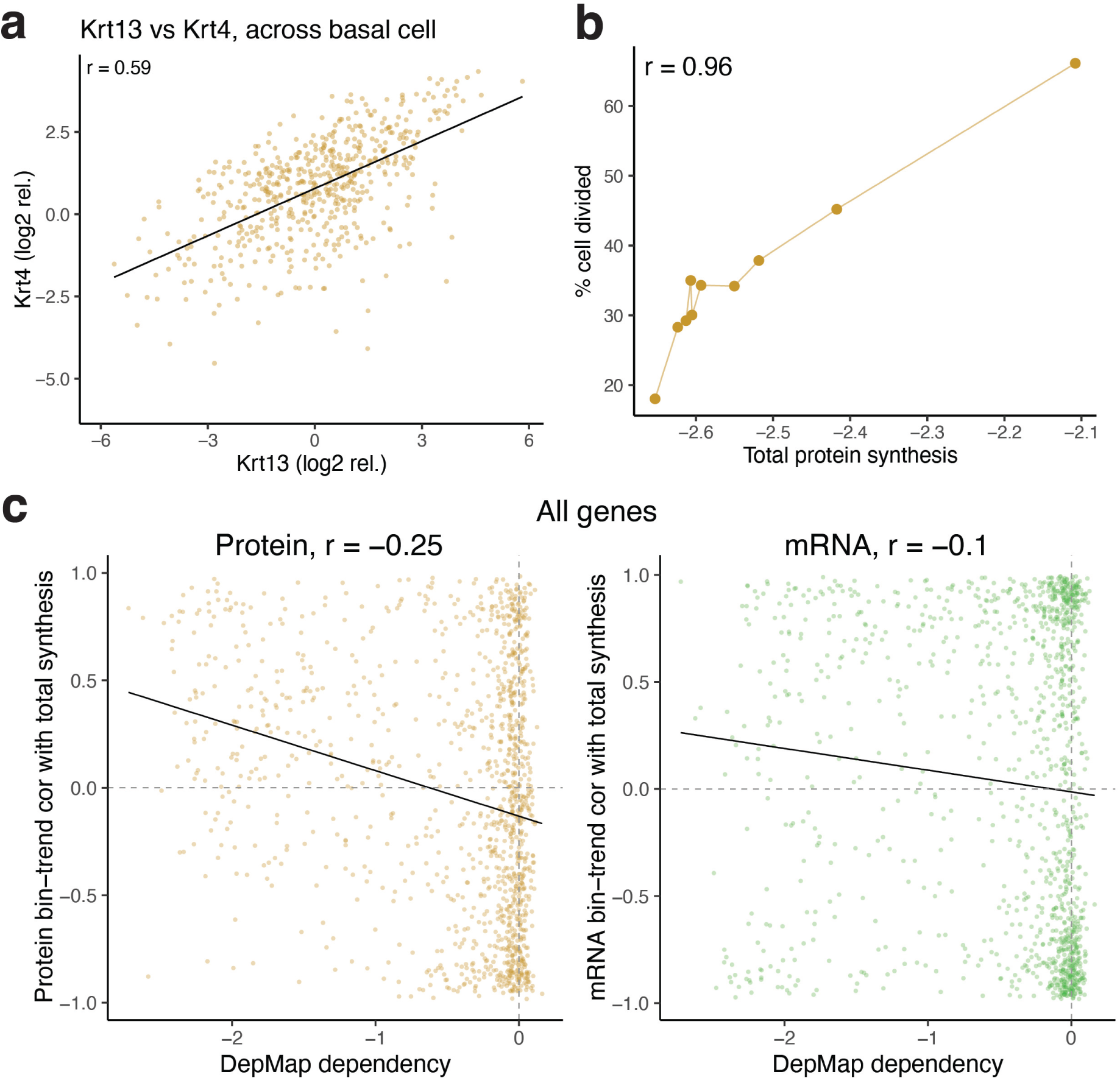
Within cell type validation figure. **a**, Scatter of Krt13 and Krt4 relative protein abundance across single basal cells. The two hillock markers co-vary strongly, Pearson *r* = 0.59, defining a within-population axis of proliferative variation, Fig. 3. **b**, Percent of basal cells that have divided during the pulse, computed within each of the 10 shared-PC1 bins, plotted against the bin-averaged total protein synthesis. **c**, Genome-wide comparison of DepMap CRISPR proliferation-dependency score against the per-gene bin-trend correlation with total protein synthesis, for protein-level (left) and mRNA-level (right) trends.

**Supplemental Fig. S2.**
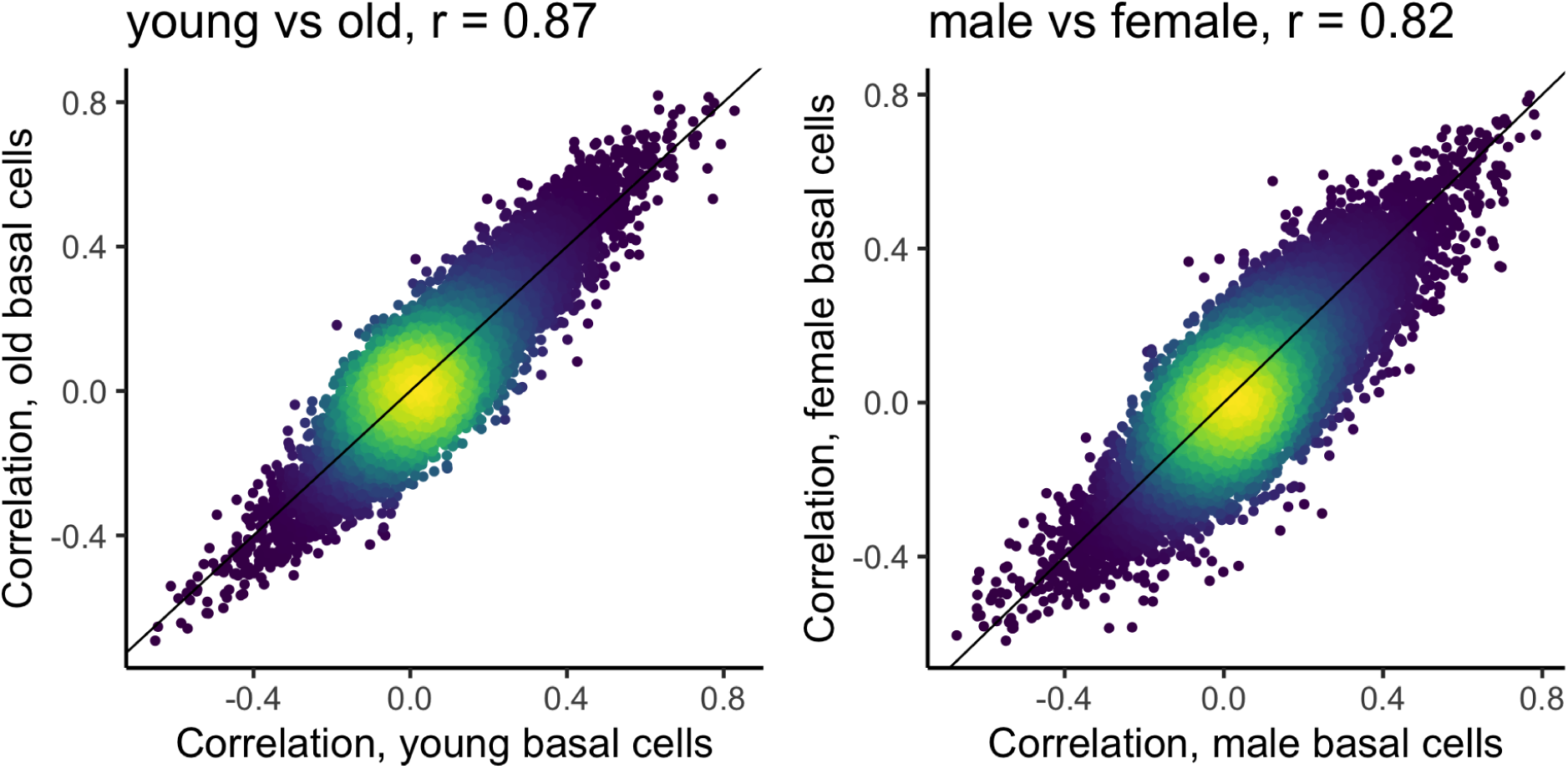
Age and sex effect on covariation structure. Pairwise protein-protein correlations were computed independently in the two groups, young and old or male and female mice. Pairs with fewer than 100 pairwise observations.

**Supplemental Fig. S3.**
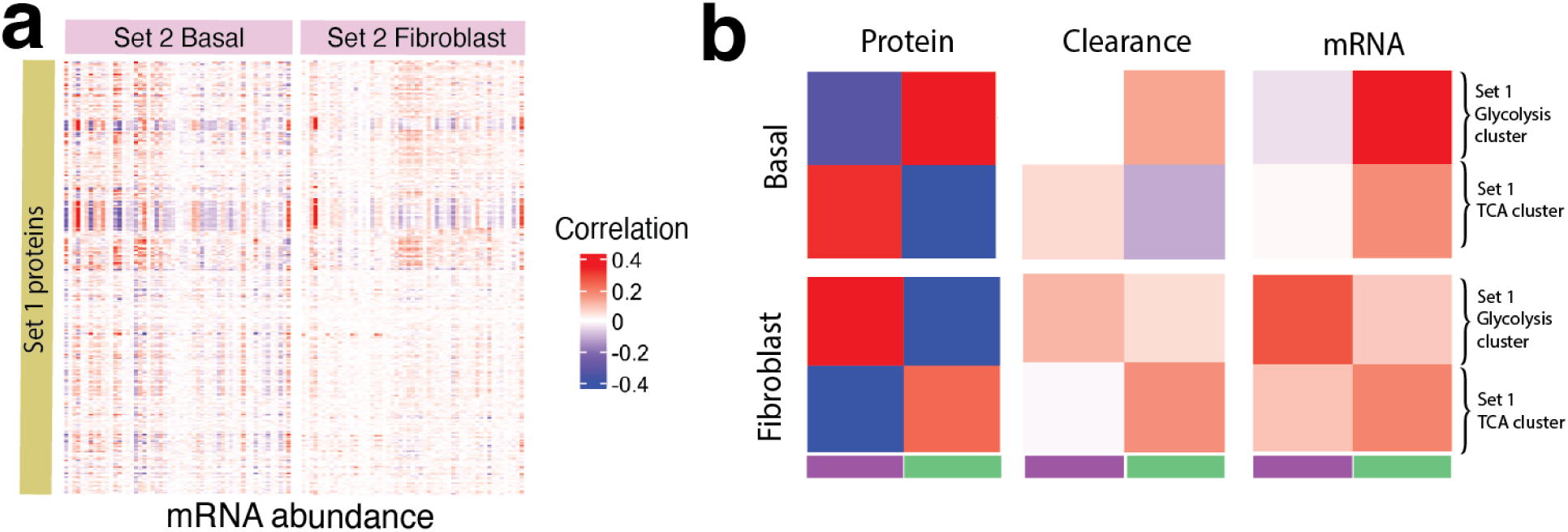
Analyzing the differences between mRNA and protein correlation vectors. **a**, mRNA-mRNA correlations corresponding to the protein-protein correlations displayed in Fig. 6d. **b**, The protein abundance, protein clearance, or mRNA abundance for gene products from the clusters in Fig. 6d were averaged for each cluster, and the cluster averages correlated. The protein clusters have the largest correlations that exhibit qualitative similarity with both clearances and mRNA correlations, suggesting contributions of these processes to the observed cell-type specific changes in protein covariation.

